# Genome instability drives epistatic adaptation in the human pathogen *Leishmania*

**DOI:** 10.1101/2021.06.15.448517

**Authors:** Giovanni Bussotti, Laura Piel, Pascale Pescher, Malgorzata A. Domagalska, K. Shanmugha Rajan, Smadar Cohen-Chalamish, Tirza Doniger, Disha-Gajanan Hiregange, Peter J Myler, Ron Unger, Shulamit Michaeli, Gerald F. Späth

## Abstract

How genome instability is harnessed for fitness gain despite its potential deleterious effects is largely elusive. An ideal system to address this important open question is provided by the protozoan pathogen *Leishmania*, which exploits frequent variations in chromosome and gene copy number to regulate expression levels. Using ecological genomics and experimental evolution approaches we provide first evidence that *Leishmania* adaptation relies on epistatic interactions between functionally associated gene copy number variations in pathways driving fitness gain in a given environment. We further uncover post-transcriptional regulation as a key mechanism that compensates for deleterious gene dosage effects and provides phenotypic robustness to genetically heterogenous parasite populations. Finally, we correlate dynamic variations in snoRNA gene dosage with changes in rRNA 2’-*O*-methylation and pseudouridylation, suggesting translational control is an additional layer of parasite adaptation. *Leishmania* genome instability is thus harnessed for fitness gain by genome-dependent variations in gene expression, and genome-independent, compensatory mechanisms. This allows for polyclonal adaptation and maintenance of genetic heterogeneity despite strong selective pressure. The epistatic adaptation described here needs to be considered in *Leishmania* epidemiology and biomarker discovery, and may be relevant to other fast evolving, eukaryotic cells that exploit genome instability for adaptation, such as fungal pathogens or cancer.

**One Sentence Summary:** Epistatic interactions harness genome instability for *Leishmania* fitness gain.

## Main Text

Darwinian evolution plays a central, yet poorly understood, role in human disease. Iterative rounds of genetic mutation and environmental selection drive tumor development, microbial fitness and therapeutic failure. Genome instability is a key source for genetic and phenotypic diversity, often defining disease outcome (*1–4*). However, the mechanism(s) by which genome instability is harnessed for fitness gain despite its potential deleterious effects remain largely elusive. Here we investigate this question in the protozoan parasite *Leishmania*, which causes devastating human infections. These parasite cycle between insect-stage promastigotes that infected *Phlebotomus* sand flies, and intracellular amastigotes that proliferate inside phagolysosomes of mammalian macrophages. Genome instability is hallmark of *Leishmania* biology, since these parasites lack promoter-dependent gene regulation (*5, 6*), but exploit chromosome and gene copy number variations to regulate mRNA abundance by gene dosage (*7–11*). In the absence of confounding transcriptional control, *Leishmania* thus represents an ideal system to investigate the role of genome instability in fast-evolving eukaryotic cells.

Here we uncover complex epistatic interactions between gene copy number variations and compensatory, transcriptomic responses as key processes that harness genome instability for fitness gain in *Leishmania*. Our data may be broadly applicable to pathogenic fungi or cancer cells, known to exploit genome instability for adaptation.

## Results

### *Leishmania* genomic adaptation is governed by epistatic interactions between gene copy number variations

We first assessed the level of copy number variation across 204 *Leishmania donovani* clinical isolates from the Indian sub-continent (ISC) (*12*). This collection includes a core group of 191 strains that are genetically highly homogenous as judged by the small number of single nucleotide variants (SNVs) (<2,500 total), which provided us with a useful benchmark to study the dynamics of copy number variations (CNVs) across a large number of quasi-clonal populations. DNA read depth analysis of these isolates revealed important CNVs in both coding and non-coding regions, with amplifications and deletions affecting respectively 14% and 4% of the genome (**Fig. 1A-B, Tables S1-S2**). Analysing the statistical association of observed CNVs with repetitive sequence elements uncovered 10 novel DNA repeats that could drive *Leishmania* genome instability through microhomology-mediated, break-induced replication as observed for human CNVs (*13*) (**Fig. 1C**, **Fig. S1, Table S3**). Gene dosage changes were not random but clearly under selection as judged by the reproducibility of genetic interactions across independent isolates and the enrichment of amplified genes in biological functions associated with fitness gain. Statistically significant interactions were observed between positive (correlating) and negative (anti-correlating) read depth variations (**Fig. 1D**, **Fig. S2-S3**, **Table S4-S5**), including a highly connected Network Cluster (NC) containing 60 co-amplified tRNA genes encoded on 16 different chromosomes (NC1, **Fig. 1D**, **Fig. S4**, **Tables S6-S7**). Natural selection of gene CNVs is further supported by (i) their independent emergence across phylogenetically distinct strains providing evidence for evolutionary convergence, (ii) the very high copy number observed for certain genes (up to 30-fold) suggesting strong, positive selection, and (iii) the global enrichment of read depth variations in phenotypically silent, intergenic regions, suggesting purifying selection against deleterious effects caused by gene CNVs (**Fig. 1E-G**, **Fig. S5, Table S8-S10**). Together our data provide the first evidence that *Leishmania* genomic adaptation is governed by gene CNVs through highly dynamic, functional interactions that are under natural selection. These interactions define a novel form of epistasis at the gene (rather than nucleotide) level, with the phenotypic effect of a given gene amplification being dependent on co-amplification of functionally related genes.

**Fig. 1.**
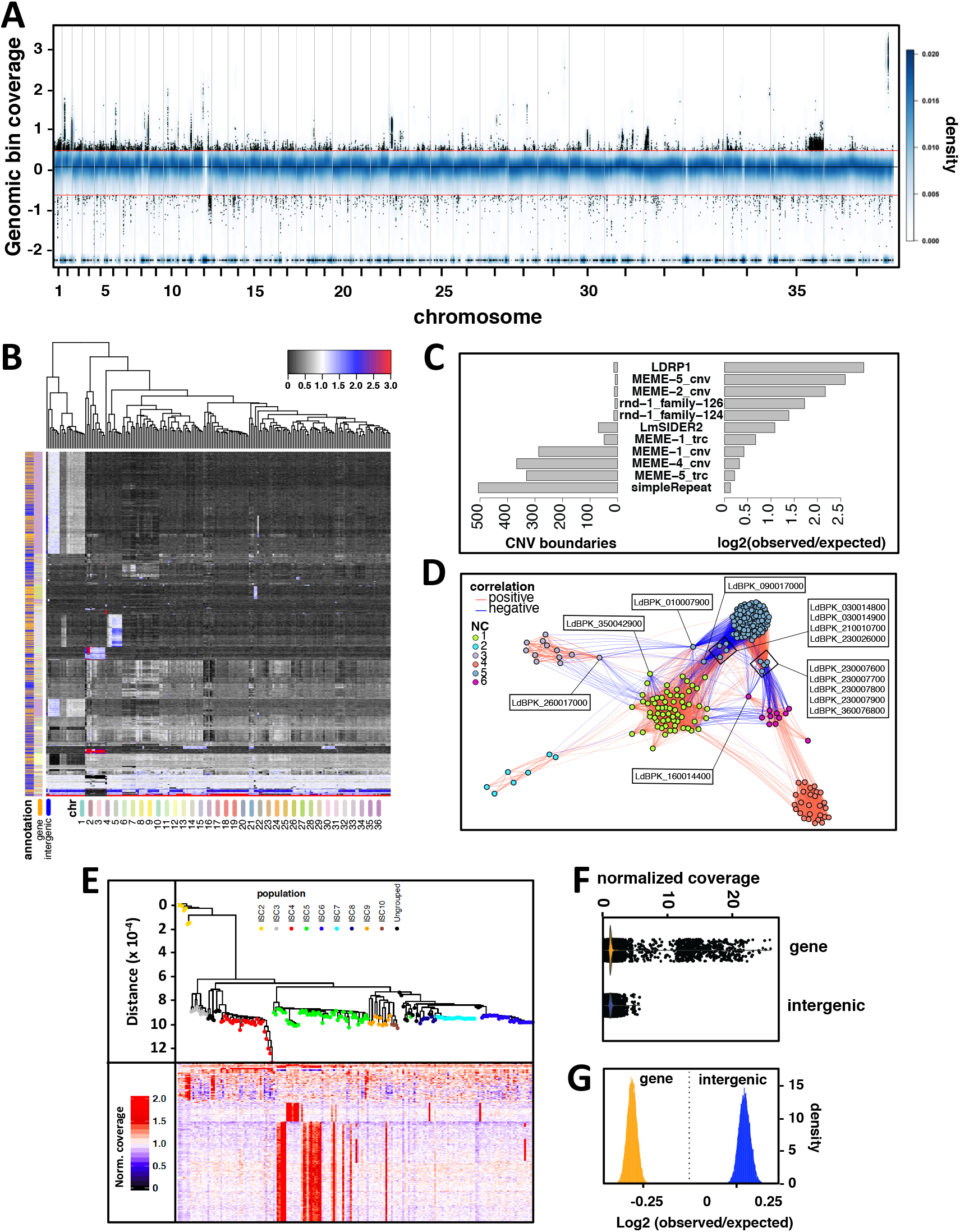
Genome-wide mapping of copy number variations (CNVs), their environmental selection, and epistatic interactions. (**A**) Genome-wide normalized coverage values in natural logarithm scale (*y*-axis) across the 36 chromosomes (*x*-axis) for 204 *Leishmania donovani* field isolates from the Indian Sub-continent (*12*). The *x*-axis reports the position of genomic windows along the chromosomes. The smoothed blue color represents the 2D kernel density estimate of genomic bins. A sample of 50,000 genomic bins with normalized coverage ≥ 1.5 or ≤ 0.5 are materialized as black dots. The black horizontal line and the two red lines indicate normalized coverage values of 1.5, 1 and 0.5. (**B**) Heatmap generated for the 204 clinical isolates (columns) showing their phylogenetic relationship as a function of CNV bins (rows). The color scale reflects the deviation from the minimum bin coverage observed across all genomes. The presence of annotated genes and the chromosomal location of the CNVs are indicated by the two colored columns on the left. The color code is defined in the legend below the plot. (**C**) Association between CNVs and repetitive elements. The bar plots show the number of observed overlap instances between the boundaries of the CNV regions and repetitive elements (left panel), and the log2 ratio between the observed and expected overlap events over 10,000 randomizations (right panel). (**D**) Gene CNV network analysis. The nodes represent gene CNVs while the edges indicate statistically significant positive (red) and negative (blue) correlations observed in the 204 field isolates. The nodes are colored according to the predicted network clusters (NC). (**E**) Phylogenetic tree based on SNVs (>90% frequency) (upper panel) for the ISC core population comprising 191 isolates (*12*) (not including the distant ISC1 strains (*38*) – for full phylogeny see **Fig. S5**). The heatmap (lower panel) shows the normalized gene sequencing coverage across genomes (rows). To ease the visualization, gene amplifications with normalized coverage > 2 are indicated as 2. (**F**) Violin plot showing the distributions of the normalized genomic coverage values of the collapsed CNV positions (dots) matching genic and intergenic regions. (**G**) Log2 ratio distributions of observed and expected nucleotide overlap between collapsed CNV regions and gene/intergenic annotations.

### Epistatic adaptation drives polyclonal fitness gain *in vitro*

We next used an experimental evolution approach to directly assess the link between epistatic interactions and fitness gain in hamster-derived *L. donovani* parasites during adaptation to *in vitro* culture (*10*). Following normalization for karyotypic variations (**Fig. S6A, Table S11**), changes in read depth were monitored between passages 2 (P2, two weeks in culture) and 135 (P135, 36 weeks in culture), corresponding to approximately 20 and 3,800 generations. Our analysis revealed co-amplification of coding and non-coding (nc) genes and gene clusters that are functionally linked to fitness gain in culture (*i.e.* accelerated cell proliferation), including genes encoding for rRNAs, tRNAs, snRNAs, snoRNAs, SLRNAs, and ribosomal proteins (**Fig. 2A**, **Fig. S6B** and **Tables S12-S14**). Epistatic adaptation thus increases translation efficiency, thus overcoming the major rate limiting steps for fast growing populations (*14*). This functional link between a given fitness phenotype and its underlying epistatic network opens unexplored venues for the discovery of *Leishmania* biomarkers, which may be represented by complex genetic interactions rather than individual loci.

**Fig. 2.**
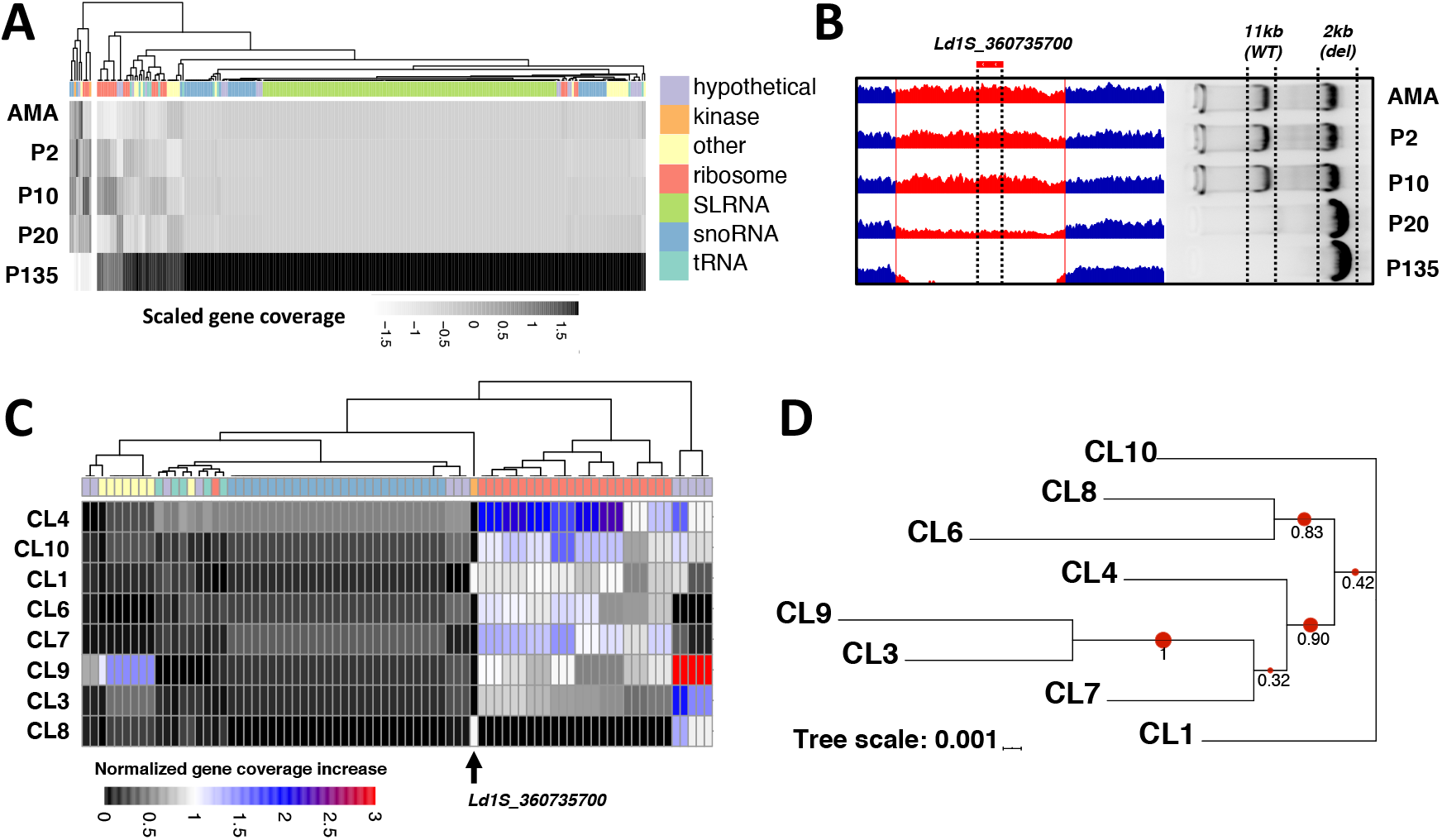
Longitudinal analysis of gene CNVs during fitness gain in culture. (**A**) Heatmap generated by plotting gene read depth values (columns) across *L. donovani* amastigotes isolated from infected hamster spleen (AMA) and derived promastigotes evolving in continuous culture for 2, 10, 20 and 135 passages (P2, P10, P20, P135) (rows). The gray level reflects the scaled normalized gene coverage as indicated in the figure. (**B**) Screenshot of the IGV genome browser showing gradual loss of the NIMA-like kinase gene Ld1S_360735700 during culture adaptation between splenic amastigotes (AMA), and derived promastigotes at passages P2 and P135. The right panel shows the gel-electrophoretic analysis of PCR fragments obtained from lesion amastigotes (AMA) and derived promastigotes at the indicated culture passages that are diagnostic for the WT (11kb) and the deleted NIMA-like kinase locus (2kb). (**C**) Heatmap generated by plotting gene read depth variation (columns) across eight clones isolated from the P20 population (rows). The color code is defined in the legend and corresponds to the deviation from the minimum sequencing coverage measured for that gene in all clones. The colored ribbon indicates the simplified gene annotation as shown in the legend of panel A. The deletion of the NIMA-like kinase Ld1S_360735700 is indicated by the arrowhead. (**D**) Analysis of the phylogenetic relation of the P20 clones. The red dots and numbers indicate the node bootstrap support (1 = 100%). The genetic distance is indicated by the branch length and scaled as indicated in the figure.

Unexpectedly, as well as gene amplification, we discovered that gene depletion is also a major driver for environmental adaptation and fitness gain. We identified a genomic deletion of 11 kb containing a single gene encoding for a NIMA-related kinase gene (Ld1S_360735700), which is gradually selected in the adapting promastigote population from a pre-existing mutant that was detected in splenic amastigotes (**Fig. 2B**, **Fig. S7A** and **B**). Clonal analysis of the P20 population revealed the presence of the spontaneous knockout (spo-KO) in six out of eight individual clones (**Fig. 2C**). These spo-KO clones were clearly not the descendants of a single founder cell but were of independent, polyclonal origin as judged by their distinct gene CNV profiles (**Fig. 2C**, **Table S15**), and their polyphyletic clustering based on SNVs compared to wild-type (WT) clones (**Fig. 2D, Table S16**). This evolutionary convergence strongly supports natural selection of the deletion during culture adaptation and suggests a potential role of the deleted NIMA kinase in growth restriction, which we further assessed by gene editing.

### Toxic gene dosage effects are compensated at post-transcriptional levels

Unlike the spo-KO clones, CRISPR/Cas9-generated, NIMA knock out mutants (cri-KO) (**Fig. S7C** and **D**) were not viable, while heterozygous mutants showed a strong growth defect, which was partially rescued by episomal over-expression of the NIMA kinase gene (**Fig. 3A**). This paradoxical result suggests that spo-KO cells must have evolved mechanisms that can compensate for the loss of this essential gene. Read depth analyses of the spo-KO and WT clones ruled out genetic compensation **(Table S15**). In contrast, RNAseq analysis revealed highly reproducible, compensatory transcript profiles in the six independent spo-KO clones (**Fig. 3B**, **Table S17**). Our analysis revealed reduced stability in spo-KO clones of 23 transcripts implicated in flagellar biogenesis (**Fig. S8**), which correlated with reduced motility (data not shown). Loss of motility may represent a fitness trade-off, providing energy required for accelerated *in vitro* growth. On the other hand, we identified 350 transcripts with increased abundance (**Table S17**, second sheet), which either result from increased gene dosage or mRNA post-transcriptional stabilization. Direct comparison of DNA and RNA reads depth variations allowed us to distinguish between these two possibilities and identified a set of transcripts whose expression changes between WT and spo-KO did not correlate with gene dosage. In absence of transcriptional regulation in *Leishmania* (*5*), the abundance of these transcripts is likely regulated by differential mRNA turn-over. Increased abundance was observed in spo-KO clones for the mRNA of another NIMA-related kinase (Ld1S_360735800) encoded adjacent to the deleted region, suggesting a direct, post-transcriptional compensation of kinase functions (**Fig. 3C, Table S18**). Likewise, we observed increased stability for functionally related small, non-coding RNAs, including 43 snoRNAs, and various rRNAs and tRNAs (**Fig. 3D, Table S18**), suggesting ribosomal biogenesis and translational regulation as yet another level of non-genomic adaptation. Together our data identify gene deletion and compensatory, post-transcriptional responses as novel drivers of *Leishmania* fitness gain.

**Fig. 3.**
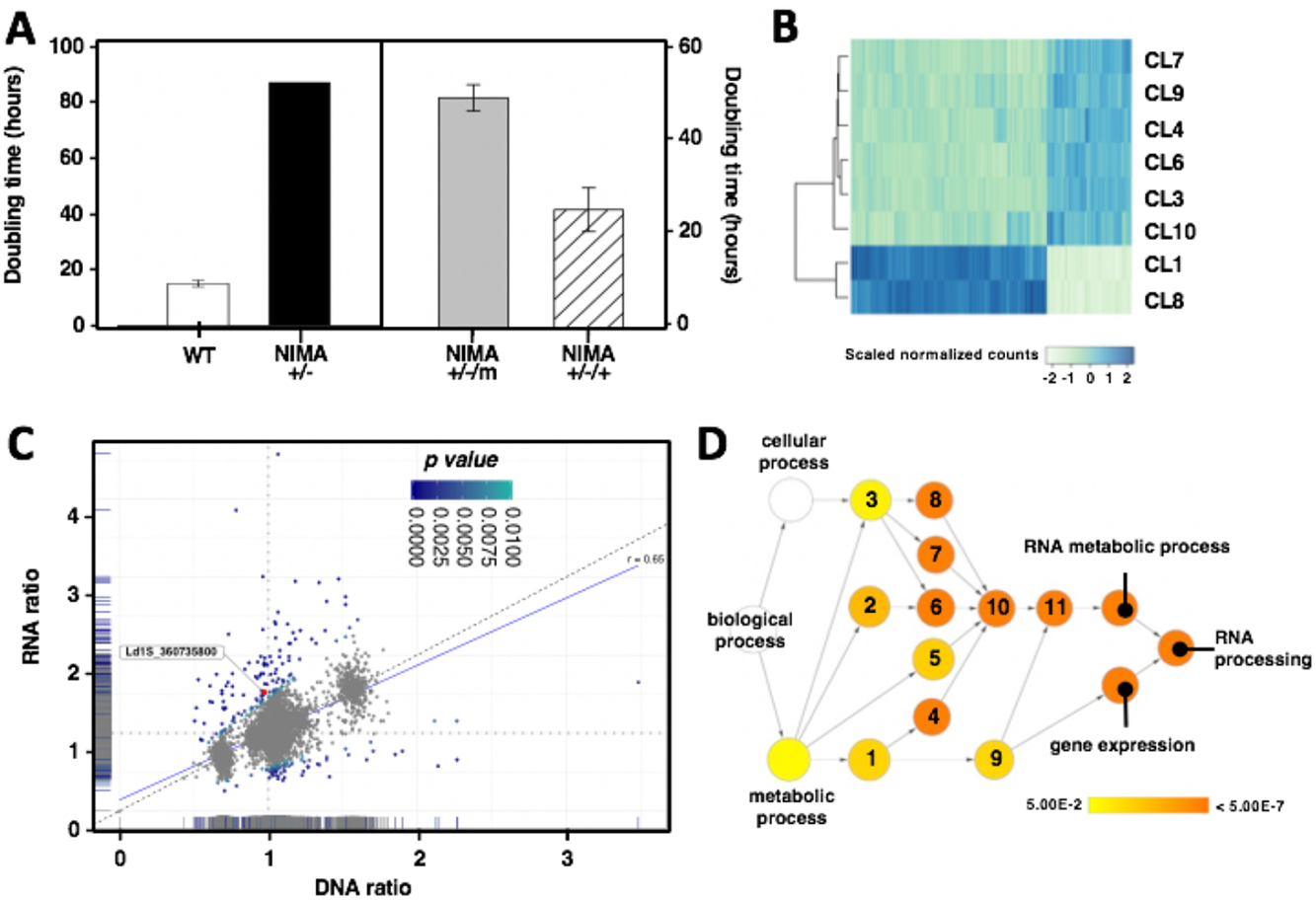
Genetic and transcriptomic analyses of the NIMA-like kinase null mutant. (**A**) Growth analysis. WT and heterozygous NIMA +/-mutants generated by CRISPR/Cas9 gene editing (left panel). Transgenic NIMA+/-parasites transfected with empty vector (NIMA+/-/m) or vector encoding for the NIMA-like kinase gene (NIMA+/-/+) (right panel). The doubling time of parasites in logarithmic culture phase is shown. (**B**) Heatmap of the scaled normalized RNAseq counts of the genes differentially expressed in the P20 clones 1 and 8 (WT) with respect to the spontaneous NIMA null mutant (spo-KO) clones 3, 6, 4, 7, 9, 10. A log2 fold change > 0.5 with adjusted p-value < 0.01 was considered significant. Darker levels of blue reflect higher expression levels as indicated by the legend. (**C**) Double-ratio scatter plot. The plot represents the ratio of the mean DNA (*x*-axis) and RNA (*y*-axis) sequencing read counts between the clones that lost the NIMA-kinase gene (CL3-4-6-7-9-10) and the NIMA-kinase wild-type clones (CL1 and CL8). Each dot represents an individual gene. The marginal distributions for DNA and RNA ratio values are displayed along the *x*- and *y*-axes. The color indicates the statistical significance level of the genes’ double-ratio scores (*i.e.* RNA ratio divided by DNA ratio) as indicated in the legend. The NIMA- like kinase homolog Ld1S_360735800 is labeled in red. The vertical and horizontal dotted lines indicate DNA and RNA ratio values of 1, while the diagonal dashed line specifies the bisector. The blue line represents a linear regression model built on the DNA and RNA ratio values and measuring a Pearson correlation value of 0.65. (**D**) Functional enrichment analysis of the biological process Gene Ontology (GO) terms for all genes in panel C showing a statistically significant double-ratio score. The node color mapping is ranging from yellow to dark orange to represent increasing significance levels, or lower adjusted p-values. White nodes are not significant. 1: organic substance metabolic process, 2: nitrogen compound metabolic process, 3: cellular metabolic process, 4: organic cyclic compound metabolic process, 5: primary metabolic process, 6: cellular nitrogen compound metabolic process, 7: heterocycle metabolic process, 8: cellular aromatic compound metabolic process, 9: macromolecule metabolic process, 10: nucleobase-containing compound metabolic process, 11: nucleic acid metabolic process.

### The non-coding RNome as a novel driver of *Leishmania* fitness gain

The selective stabilization of snoRNAs during early culture adaption (P2-P20) suggests these non-coding RNAs are key drivers in *Leishmania* fitness gain. We confirmed this possibility in long-term adapted parasites that were continuously cultured for ∼3,800 generations (P135). Read depth analysis revealed selective amplification of snoRNAs, which was confirmed in an independent, long-term evolutionary experiment conducted until passage 125 (P125, **Fig. 4A**, **Fig. S9** and **Table S19**). Surprisingly, rather than amplification of individual snoRNA genes, increased read depth was caused by the recovery of a single locus on chromosome 33 containing a cluster of 15 snoRNA genes in the P135 population, but was depleted in the original amastigote population (**Fig. 4B**). The restoration of this locus between P20 and P135 is further proof that snoRNAs are under positive selection during culture adaptation. Rather than gene amplification, the recovery of this locus is more likely due to the selection of a small sub-population that penetrated the culture between P20 and P135, similar to what we observed for the NIMA kinase spo-KO. The mosaic structure of this locus is further supported by the CNVs we detected in independent amastigotes isolates (**Fig. 4C**, **Fig. S10**). Thus, while short-term adaptation in *Leishmania* is governed by control of RNA stability, long-term adaptation occurs through more cost-efficient gene dosage regulation. Increased snoRNA abundance likely satisfies a quantitative need for ribosomal biogenesis in our fast-growing cultures. However, changes in snoRNA expression can also affect the nature and quality of ribosomes (*15, 16*).

**Fig. 4:**
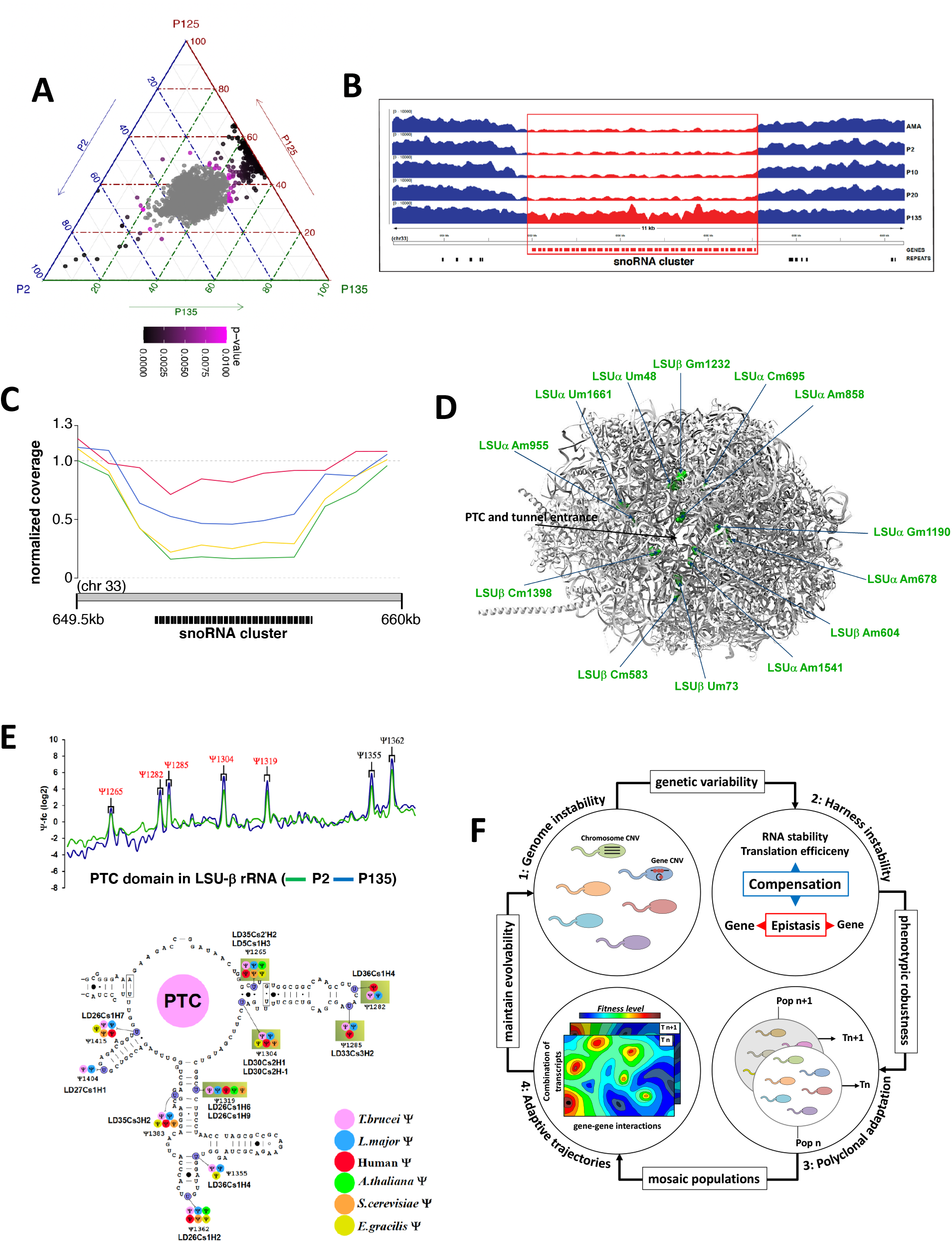
snoRNA genes are amplified in long-term adapted parasites and promote rRNA modification. (**A**) Ternary plot showing for each gene the relative abundance in culture passage P2 – P125 and P135. The axes report the fraction of the normalized gene coverage in each sample, with each given point adding up to 100. Dots with color ranging from pink to black indicate significant gene CNVs (p-value < 0.001). (**B**) Recovery of the snoRNA gene cluster CNVs during culture adaptation. The panel illustrates a genome browser representation of the sequencing depth measured in the samples at the indicated passage. Gene annotations and the predicted repetitive elements are indicated. (**C**) Line plot showing the normalized sequencing coverage (y-axis) over the snoRNA cluster region on chromosome 33 (x-axis) for the amastigote sample (AMA, green) and three independent amastigote isolates (AMAH154, red; AMA07142, yellow; AMA1992, blue) obtained from different hamster infections. (**D**) Hypermodified Nm sites are located around the functional domains of the ribosome. The complete stoichiometry of each Nm site was measured by RiboMeth-seq in P2 and P135 *L. donovani* strains as presented in **Table S20**. The hypermodified Nm sites are highlighted in green and their identity is indicated on the 3D structure of the *L. donovani* large subunit ribosome based on the previously deposited cryo-EM coordinates (Protein Data Bank (*39*) accession 6AZ3) (*40*). The location of the PTC and mRNA entrance tunnel are shown. (**E**) Hypermodified Ψs are present in the functional domains of the ribosome. The relative changes in Ψ level was measured using Ψ-seq and is presented in **Table S21**. Representative line graph of the fold change in rRNA pseudouridinylation level (Ψ-fc, log2) is presented for P2 (green line) and P135 (blue line) for the PTC domain in LSU-Ψ rRNA. The positions where the Ψ level is increased in four replicates are indicated in red. The location of Ψ sites in the rRNA is depicted on the *Trypanosoma brucei* secondary structure (*19*). Hypermodified sites are highlighted in boxes. The snoRNA guiding on each Ψ is indicated. The color code for each Ψ site is indicative of the organism where it was already reported. (**F**) Model of *Leishmania* polyclonal adaptation. (1) *Leishmania* intrinsic genome instability generates constant genetic variability. (2) Epistatic interactions between gene CNVs and compensatory responses at the level of RNA stability and translation efficiency eliminate toxic gene dosage effects and harness genome instability. (3) This mechanism generates phenotypic robustness while at the same time maintaining genetic variability, thus allowing for polyclonal adaptation. (4) The genetic mosaic structure allows for distinct adaptive trajectories (T) inside and in-between adapting populations, a process that conserves genetic heterogeneity and thus evolvability of the population despite constant selection.

In the following we assessed the possibility of such fitness-adapted ribosomes in *Leishmania* especially since snoRNA genes guiding 2’-*O*-methylation (Nm) and pseudouridine (ψ) modifications were extensively amplified. Amplification of snoRNAs should lead to increased modifications on sites that are accessible for modification. The mapping of Nm via RiboMeth-seq (*17*) revealed an increased level of modification for 18 sites by at least two fold (**Table S20**), while the mapping of pseudouridylation by ψ-seq (*18, 19*) showed increase for 5 ψ sites (**Table S21**) during adaptation from P2 to P135. Interestingly, the hyper-modified Nm sites are localized around the peptidyl-transferase centre (PTC) and mRNA entrance tunnel, whereas the hyper-modified ψ sites are located in the PTC itself (**Fig. 4D-E**, **Fig. S11**). Together, our data reveal a complex model of *Leishmania* fitness gain, where epistatic interactions between gene amplifications and compensatory responses at post-transcriptional levels harnesses *Leishmania* genome instability for polyclonal adaptation (**Figure 4F**).

## Discussion

A common strategy in microbial evolutionary adaptation is known as ‘bet-hedging’, where the fitness of a population in changing environments is increased by stochastic fluctuations in gene expression that are regulated at epigenetic levels (*20–24*). The protozoan pathogen *Leishmania* largely lacks transcriptional regulation, raising the question on how these parasites generate variability in transcript abundance and phenotype required for adaptation (*5, 6*). Our data uncover an alternative mechanism of bet-hedging that has evolved based on the unique biology of *Leishmania*.

First, we provide evidence that *Leishmania* genome instability may be driven by 10 new repetitive DNA elements associated with genome-wide amplifications and deletions, both of which are under positive and purifying selection. *Leishmania* thus compensates for the absence of stochastic gene regulation by the generation of stochastic gene CNVs, which are known to cause dosage-dependent changes in transcript abundance (*7–11*). Second, we reveal an important role for post-transcriptional regulation in *Leishmania* fitness gain, which (i) compensates for the deleterious deletion of a NIMA kinase by selectively stabilizing the transcripts of an orthologous kinase, (ii) allows for gene dosage-independent increase in the abundance of non-coding RNAs (SLRNAs, snoRNAs) required for proliferation, and (iii) provides phenotypic robustness to genetically heterogenous populations as documented by the converging transcript profiles of independent NIMA spo-KO mutants. This adaptation process guards against toxic gene dosage effects and simultaneously increases the phenotypic landscape available to *Leishmania* for adaptation *via* gene deletions and compensatory transcriptional responses. Significantly, the NIMA orthologue, as well as the SLRNA and snoRNA loci are amplified during long-term culture (**Fig. 4B**, **Fig. S6B**, **Fig. S9**), revealing a two-step adaption process reminiscent to yeast (*25*), implicating a first, post-transcriptional mechanism *via* transient changes in RNA stability, followed by a second, genomic mechanism *via* selection for stable CNVs.

The dynamic changes in snoRNA stability and gene copy number observed in our experimental evolution system identifies this class of ncRNAs as an unexpected driver of *Leishmania* fitness gain. snoRNAs guide rRNA modification and processing, as well as modification of snRNAs and additional non-coding ncRNAs (*26, 27*). Changes in snoRNA gene copy number and transcript abundance during parasite adaptation correlated with hyper-modification of rRNAs, known to affect the quality and translation specificity of the ribosome. It is interesting to speculate that these specialized ribosomes represent an additional layer of regulation at translational level that can further counteract toxic gene dosage effects, provide phenotypic robustness and adapt the proteome profile to a given environment, much like it was observed in cancer cells or during differentiation of stem cells (*28, 29*).

Our findings have important clinical implications for *Leishmania* infection. *Leishmania* adapts to various environmental cues, notably the presence of anti-leishmanial drugs. In contrast to high frequency amplifications observed during experimental drug treatment in culture (*30–32*), treatment failure and drug resistance observed during natural infection may evolve through multi-locus, epistatic interactions such as those described here, which can balance the fitness trade-off between drug resistance and infectivity (*33*). Therefore, our data define biological networks, rather than individual genes, as novel biomarkers with potential diagnostic or prognostic value. Conceivably, the epistatic mechanisms we uncover in *Leishmania* can be of broader relevance to other human pathologies caused by fast evolving eukaryotic cells exploiting genome instability for polyclonal adaptation, such as cancer cells. While single nucleotide, epistatic interactions are recognized as important drivers of tumor development (*34, 35*), the role of epistatic interactions between structural mutations and between the genome and transcriptome in drug-resistant cancer cells remains to be elucidated (*36*). In conclusion, our results propose a novel model of *Leishmania* fitness gain (**Fig. 4F**), where polyclonal adaptation of mosaic populations is driven by epistatic interactions that (i) buffer the detrimental effects of genome instability, (ii) coordinate expression of functionally related genes, and (iii) generate beneficial phenotypes for adaptation. This mechanism of fitness gain avoids genetic death by maintaining heterogeneity in competing parasite populations under environmental selection and may be generally applicable to other eukaryotic systems that adapt through genome instability.

## Acknowledgments

**Funding:** This study was supported by a seeding grant from the Institut Pasteur International Department to the LeiSHield Consortium, the EU H2020 project LeiSHield-MATI - REP-778298-1, the Fondation pour la Recherche Médicale (grant number FDT201805005619), the Flemish Ministry of Science and Innovation (MADLEI, SOFI Grant 754204) and a grant from CAMPUS France and the Israeli Ministry of Science and Technology PHC MAIMONIDE 2018-2019-Projet N° 41131ZD. We thank Cedric Notredame and Jean-Claude Dujardin for critical reading of the manuscript.

**Author contributions:** Conceptualization, G.B., L.P., P.P., M.A.D., T.D., R.U. S.M. and G.F.S.; Methodology, G.B., L.P., P.P., K.S.R, S.C.C, D.G.H., T.D., R.U. and G.F.S.; Software, G.B., T.D. and R.U.; Formal Analysis, G.B., L.P., P.P. and G.F.S.; Investigation, G.B., L.P., P.P. and G.F.S.; Writing – Original Draft, G.B., L.P., P.P. and G.F.S.; Writing – Review & Editing, G.B., L.P., P.P., M.A.D., T.D., P.J.M, R.U. S.M. and G.F.S.; Supervision, G.F.S., R.U. and S.M.; Funding Acquisition, G.F.S., S.M.

**Competing interests:** Authors declare no competing interests.

## Data and materials availability

Reads were deposited in the Sequence Read Archive (SRA) database (*37*) and are publicly available under accession no PRJNA605972. All data is available in the main text or the supplementary materials.

## Supplementary Materials

### Materials and Methods

#### *L. donovani* strains, culture conditions and cell cloning

Culture-adapted *L. donovani* field isolates maintained for more than 20 *in vitro* passages (P) from the Indian Sub-Continent (ISC) and previously subjected to whole genome sequencing analysis were used in our study (*12*). *Leishmania (L.) donovani* strain 1S2D (MHOM/SD/62/1S-CL2D) (Ld1S) was obtained from Henry Murray, Weill Cornell Medical College, New York, USA and maintained by serial passages in hamsters. Amastigotes were recovered from infected hamster spleen and differentiated into promastigotes in M199 medium supplemented with 10% FCS, 25mM HEPES pH 6.9, 4mM NaHCO3, 1 mM glutamine, 1x RPMI 1640 vitamin mix, 0.2 µM folic acid, 100 µM adenine, 7.6 mM hemin, 8 µM biopterin, 50 U ml^-1^ of penicillin and 50 µg ml^-1^ of streptomycin. Promastigotes were serially passaged once stationary phase was reached for either 2, 10, 20, or 135 passages corresponding to respectively 20, 60, 190, and 3,800 generations. An independent *L. donovani* isolate was grown in culture for over 33 months until P125 (corresponding to approximately 3,500 generations). Three independent splenic amastigotes populations were recovered from as many infected hamsters using the same culture conditions. Serial dilutions of passage 20 (P20) promastigotes were plated on M199 Agar plates and 8 clones were recovered and expanded for 2 passages in liquid culture (*10*).

#### Nucleic acid extraction and deep sequencing

All sequencing analyses were performed using Illumina short-read technology. DNA extraction and sequencing protocols of the *L. donovani* field isolates from the Indian Sub-Continent (ISC) are described in (*12*) DNA extraction and sequencing analysis of *L. donovani* LD1S amastigotes (AMA) and derived promastigotes at passages P2, 10, and 20 are described in (*10*). For new amastigote isolates (AMAH154, AMA1992, AMA07142) and promastigote strains (P125, P135), DNA extractions were performed using DNeasy blood and tissue kits from Qiagen according to the manufacturer’s instructions. Nucleic acid concentrations were measured and the DNA quality was evaluated by measuring the OD ratios 260/280 and 260/230 and by agarose gel-electrophoresis. Between 2 to 5µg of DNA were used for sequencing.

For samples AMAH154, AMA1992, AMA07142, P125 and P135, short-insert, paired-end libraries were prepared with the KAPA Hyper Prep Kit (Kapa Biosystems). The libraries were then sequenced using TruSeq SBS kit v3-HS (Illumina Inc., CA, USA). Multiplex sequencing was performed using HiSeq 2000 flowcell v3, generating 2×101bp paired-end reads, according to standard Illumina procedures. Reads were deposited in the Sequence Read Archive (SRA) database (*37*) and are publicly available under accession no PRJNA605972.

#### Whole genome sequencing data analysis

Genomic reads were aligned and the nucleotide sequencing depth was measured and normalized as previously described (*41*). For the genome-wide detection of CNVs across the 204 ISC isolates, the LdBPKv2 reference genome was partitioned into adjoining intervals of a fixed length of 300bp (bins), and the depth of coverage of each bin was measured and compared across samples. The mean sequencing coverage normalized by the median chromosome coverage was computed for each bin as already described (*41*). In **Fig. 1A** and **1B** and to compute the genome fraction of amplified or depleted regions, all maxi-circle DNA bins as well as the bins with a median normalized coverage > 0.01 but mean read mapping quality score (MAPQ) < 50 were discarded. We referred to genomic bins as being deleted, depleted or amplified respectively for normalized sequencing coverage values ≤ 0.01, < 0.5 and > 1.5. **Fig. 1B** shows the copy number variant bins across the samples. To account for differences in ploidy levels, the bin copy number was evaluated as the bin sequencing coverage normalized by the chromosome median sequencing coverage (i.e. copy number per haploid genome). Bins showing a copy number variation of at least 1 across the 204 ISC isolates were considered in the analysis. The heatmap color scale reflects the difference in bin normalized read depth with respect to the minimum value measured across all genomes. Black indicates no variation. White, blue and red correspond to increases in normalized coverage levels of respectively 1, 2 and 3. Coverage variations greater than 3 were down sized to 3. The analyses shown in **Fig. 1F** and **1G** rely on CNV regions potentially spanning multiple genomic bins aggregated into individual larger units. We refer to such grouped bins as “collapsed bins”. Groups of genomic CNV bins separated by less than 1,200 bases were merged together, and their normalized sequencing coverage averaged. To account for potential unannotated UTR regions in **Fig. 1F** and **1G**, CNV regions were considered intergenic if located at a distance of at least 100bp from annotated genes. The line plot in **Fig. 4C** illustrates the sequencing coverage in 1,000bp long bins normalized by chromosome 33 median sequencing coverage.

Likewise, sequencing coverage was used to estimate and compare gene copy numbers. The mean sequencing coverage of each gene was estimated and normalized as previously published (*41*). Reads with a mean MAPQ score < 50 were filtered because not directly quantifiable with the sequencing technology we used. Genes with a variation of normalized coverage > 1 in the 204 field isolates were considered to be copy number variant. The heatmap in **Fig. 1E** includes the genes (rows) showing convergent amplification (normalized coverage > 1.5) across 191 ISC core group (*12*) isolates (columns). Genes amplified only in isolates sharing a common ancestor were excluded from the analysis. **Fig. 2A, 2C, 4A** and **S9** show gene copy number variations across different data sets. Genes supported by reads with and average MAPQ score < 50 were grouped into clusters based on sequence similarity, quantified as cluster entities and evaluated for possible variation across samples (gene cluster CNVs). Briefly, the stranded sequences of low MAPQ genes were grouped with cd-hit-est (version 4.6.8) (*42*) with options “-s 0.9 -c 0.9 -r 0 -d 0 -g 1” selecting clusters of highly similar genes. Samtools view (version 1.3) and BEDTools coverage (version 2.25.0) were used to measure the mean sequencing depth of the individual low MAPQ genes and were run, respectively, with options “-F 1028” and “-d -split”. For the mean coverage estimate possible intragenic gap regions were not considered. The mean coverage of each gene was normalized by the median coverage of its chromosome. Eventually, the normalized mean coverage of all genes in a cluster was averaged to compute the cluster support. **Fig. 2A** shows the gene and cluster increasing in copy number between amastigote (AMA) and P135, using as a cutoff an increment in normalized coverage of at least 0.5. **Fig. 2C** represents the gene and cluster CNV detected in the clones using as a cutoff a minimum variation in normalized coverage of at least 0.5. Black indicates no variation. White, blue and red reflect increasing values in normalized coverage levels of respectively 1, 2 and 3. Coverage variations greater than 3 were down sized to 3. For the analysis illustrated in **Fig. 4A** the coverage of each gene was evaluated as the number of mapping reads excluding duplicates, increased by a pseudo-count of 1, and normalized by the median number of reads per gene in the respective samples. In **Fig. S9** the mean gene sequencing coverage values were normalized by the median of the mean gene coverage values in the respective P125 and P135 samples. The genes with mean MAPQ < 50 in either sample were discarded.

To represent the sequencing coverage of selected genes and genomic areas we used the Integrative Genomics Viewer (IGV) (*43*). In **Fig. 2B**, **4B**, **S6B** and **S7A** the sequencing coverage tracks were produced with bamCoverage from the deepTools suite (*44*) (version 2.4.2) with options "--binSize 10 --smoothLength 30" and ignoring duplicated reads. Normalization of reads per kilobase per million reads (RPKM) was applied on separate chromosomes to render the coverage comparable across samples and ploidy levels.

We determined chromosome aneuploidy in our samples sets based on sequencing coverage. In **Fig. S6A** and **S10**, the normalized sequencing coverage was binned in contiguous 2,500bp windows for each sample and for each chromosome, and the distribution of the windows’ mean coverage scores were displayed. Sequenced positions where more than 50% of the reads showed a MAPQ lower than 50 were not considered in the analysis. To estimate chromosome copy number differences, the window coverage was normalized by the median coverage of the chromosome showing a stable disomic level across samples (chromosomes 21-25 respectively for **Fig. S6A** and **S10**) and multiplied by two. The distribution of the mean window coverage was compared with the R (*45*) Wilcoxon test and the p-values of the comparisons are reported in **Table S11.**

#### Reference genomes

The analyses regarding the use of the Sudanese *L. donovani* strain 1S2D (Ld1S) were performed utilizing its PacBio genome assembly available from the NCBI website under the biosample (*46*) accession n° SAMN07430226 and the bioproject (*46*) accession n° PRJNA396645. To produce Ld1S genome annotation we combined different approaches. We ran the Companion pipeline (*47*) without using transcript evidence and exploiting *L. major* Friedlin as a reference organism to specify the models for gene finding and functional annotation transfer. The predicted gene identifiers were renamed to match chromosomal localization. Then we complemented Companion annotations with LdBPKv2 (*7*) homology-based predictions. We used NCBI-tblastn (version 2.2.28) (*48*) to search the amino acid sequence of LdBPKv2 genes with an e-value threshold of 0.01. To enable the screening of non-coding sequences we also ran gmap (version 2015-07-23) with options “--npaths 30 --no-chimeras --nosplicing –prunelevel --min-trimmed-coverage 0.9 -- min-identity 0.9” searching Ld1S homologs of all LdBPKv2 genes. All the significant NCBI- tblastn and gmap predictions that were not overlapping on the same strand with Companion annotations were retained. BEDTools merge (version 2.25.0) was used to combine overlapping predictions into single gene annotations spanning all of the combined predictions. Eventually NCBI-blastn was used to search in the Ld1S intergenic space the *Leishmania major* Friedlin UsnRNA, snoRNA, SLRNA and 7SL non-coding RNA classes. Overall, the Ld1S genome encodes for 10,532 genes: 8,850 defined by the Companion pipeline, 104 defined by homology with LdBPKv2 genes, and 1,578 defined by homology with *L. major* Friedlin ncRNA classes.

Ld1S gene functions were inferred adopting a hierarchical approach. Firstly, we transferred the function of *L. donovani* strain BPK282A1 (LdBPK282A1) homolog genes available from TriTrypDB (*49*) (downloaded the 05/07/2019). To this end, OrthoFinder (*50*) (version 2.2.7) was used with the DIAMOND (*51*) search program to establish orthology relation between LdBPK282A1 (downloaded from the Sanger FTP server ftp://ftp.sanger.ac.uk/pub/project/pathogens/ on the 12/10/2018) and Ld1S gene annotations. NCBI-blastn was run with options “-dust no -soft_masking false -evalue 10” to scan and try to rescue the low complexity genes were OrthoFinder failed to determine homology. When “one-to-many” homologs occurred (i.e. one Ld1S gene matching multiple significant LdBPK282A1 homologs), the function of all individual homologs was concatenated in a single, not-redundant functional assignment. For the genes lacking functional annotation, or reported as “hypothetical proteins” we assigned the function reported in the Companion GAF output file or the transcript type if no description was available. Then, if no annotation was found other than “hypothetical proteins”, we sought to infer the gene function using HMMer (*52*) (version 3.1b2) to scan the gene protein sequences against EGGNOG kinetoplastida database (*53*) (version 4.5) (http://eggnogdb.embl.de/download/eggnog_4.5/data/kinNOG/), and ultimately combining the annotated functions in EuPathDB (*54*) and UniProt (*55*) (downloaded the 20/09/2019). To extract the UniProt function annotations we queried the Gene Ontology Annotation (GOA) (*56*) database with the Uniprot identifiers of the LdBPK282A1 homologs of Ld1S genes.

In order to assign the Gene Ontology Identifiers (GO IDs) we combined the GOA-derived identifiers with the ones available from the corresponding orthologs in target species: LdBPK, *L. infantum*, *L. major*, *L. mexicana*, *Typanosoma brucei brucei 927* (Tbru) and *Typanosoma cruzi* (Tcru). For each target species we retrieved both the “curated” and “computed” GO IDs from TriTrypDB on the 11/09/2019. OrthoFinder with the DIAMOND search program was applied to establish orthology between the genes in Ld1S and in target species. In “one-to-many” orthology relations we concatenated all the non-redundant GO IDs from all the homologs. The GO IDs were then assigned based on the hierarchy: LdBPK curated > LdBPK GOA > *L. infantum* curated > *L. major* curated > *L. mexicana* curated > Tbru curated > Tcru curated > LdBPK computed > *L. infantum* computed > *L. major* computed > *L. mexicana* computed > Tbru computed > Tcru computed. The GO IDs were assigned if not present in any higher rank GO ID data set. The GO IDs of snoRNAs, UsnRNA, SLRNA and 7SL classes defined by homology with *L. major* Friedlin genes were manually attributed. Overall, we assigned biological process (BP), molecular function (MF) and cellular component (CC) GO IDs to 5,246, 4,521 and 7,236 Ld1S genes.

All analyses regarding the 204 Indian sub-continent (ISC) isolates (*12*) used the Nepalese PacBio *L. donovani* (LdBPKv2) genome and annotation as reference (*7*). LdBPKv2 gene function was inferred adopting a hierarchical approach similar to the one described for Ld1S. Higher priority was given to TriTrypDB, followed by Companion GAF (available from (*7*)) and EGGNOG functional annotations.

#### Repeat analysis

We annotated the repetitive elements in the LdBPKv2 genome producing a comprehensive dataset (**Table S3**). To build this dataset, we merged the repetitive elements identified by different approaches in three distinct repeat sets and present with at least 5 copies in the genome.

The first set of repetitive elements accounts for the interspersed repeat elements predicted with RepeatMasker (*57*) (version 4.0.8) with options “-e crossmatch -gff -xsmall -s”, scanning the genome to identify short simple/low complexity repetitive elements and interspersed repetitive elements. RepeatMasker was run in combination with the Repbase (*58*) library to identify *Leishmania*-specific and ancestral repeats, and with the output of RepeatModeler (version 1.0.11) (*59*), a pipeline to identify transposable elements run with option “-engine ncbi”. Redundant, RepeatModeler-defined elements overlapping with Repbase elements were discarded. RepeatMasker output was processed collapsing self-inclusive low-complexity motifs to the shortest identical sub-motif (e.g. TAGTAG -> TAG) and merging together overlapping repeats elements of the same type (**Table S3**).

The second set of repetitive elements consists in the DNA motifs enriched at the boundaries of the CNV regions we detected in the core group of 191 ISC *L. donovani* field isolates, excluding the “Yeti” strains (*12*). Such boundaries were defined as the genomic regions spanning 1kb and flanking both sides of each CNVs. The CNVs defined in the core group include the 300bp genomic bins with average read MAPQ score > 50, and showing a variation of normalized sequencing coverage score > 1 in the sample set. The bins localized within 6kb were collapsed and their sequencing scores averaged. Significant DNA motifs were detected using MEME (version 5.0.4) (*60*) with option “-dna -revcomp -nmotifs 20 -evt 0.01 -mod anr” and specifying a custom background model of order 3 with fasta-get-markov, a tool included in the MEME suite. The sequences we used to build such background model were randomly extracted from the LdBPKv2 genome with BEDTools shuffle (*61*) (version 2.25.0) in order to resemble the ones scanned by MEME in terms of number and size, but not including them. The DNA motifs predicted with MEME were then mapped genome-wide with MAST using the options “-hit_list -comp -remcorr -mt 0.0000001”.

The third set of repetitive elements includes the DNA motifs enriched at the boundaries of tandem repeat cluster (TRC) regions. TRCs are characterized by the tandem repetition of large genomic segments. To detect TRCs in the LdBPKv2 genome we streamlined a pipeline including several steps and relying on several bioinformatics tools. First, we detected inexact repetitive elements on individual chromosomes. For this purpose we ran nucmer (version 3.23) (*62*) with options “--maxmatch --nosimplify”, and show-coords (version 3.23) (*62*) with options “-r -T -o -l -d -I 80 -L 50”. Possible overlapping predictions were collapsed with BEDTools merge (*61*). To remove distantly related elements the resulting elements were re-clustered by sequence similarity with cd-hit-est (version 4.6.8) with options “-d 0 -g 1”. Then to refine the element boundaries we compared all-versus-all sequences in each cluster with NCBI-blastn (version 2.2.28) (*48*) with options “-dust yes -soft_masking true -evalue 0.01”, retaining the elements longer than 50bp. Next, we processed each cluster grouping the sequences based on a maximum distance cutoff of 2kb, thus ensuring that the repetitive element occur in the same genomic area. Then the clusters found to be intersecting each other were pooled together to generate the TRCs. Eventually the TRCs were processed enforcing a minimum distance between an element and the one downstream of at least 100bp. The genomic space between the elements inside the TRC and the regions of 1kb flanking each TRC were evaluated to identify enriched DNA motifs. Significant motifs were detected with MEME and mapped genome-wide with MAST as already described for the CNV boundaries.

#### CNV association analyses

We evaluated weather LdBPKv2 CNVs are significantly associated with genes and intergenic regions utilizing GAT (*63*), a tool for testing the significance of overlap of genomic intervals. In this analysis we considered 10,000 simulations in which we assigned random positions to the collapsed bin CNV regions. For each simulation we evaluated the number of nucleotides overlapping the gene and intergenic regions, thus implementing the null models that the CNV set of intervals is placed independently. The genomic regions accessible for simulation include (i) the 36 chromosomes of the LdBPKv2 genome, excluding the assembly gaps, (ii) the genomic positions where more than 50% of the reads have a MAPQ < 50, and (iii) the regions shorter than 20kb not containing any CNV. To account for possible non-annotated UTR elements, we expanded the gene coordinates by 100bp on both sides, and consequently shrunk the intergenic regions.

GAT was also used to determine the association between the LdBPKv2 CNV boundaries and the comprehensive repeat dataset. For each of the 10,000 simulations we evaluated the number of CNV boundaries overlapping each repeat type. For this purpose, all low-complexity motifs were collectively considered as a single class named “simpleRepeats”. The CNV boundaries included a 1kb area flanking the CNV, plus the segments of 150bp spanning from the CNVs extremities and moving toward their center. Eleven repeat types were found significantly associated to CNV regions (**Fig. 1C**).

#### Network analysis

To explore the interaction between gene CNVs we performed a network analysis. In the graph shown in **Fig. 1D**, the edges indicate significant gene CNVs correlation with adjusted p-value lower than 0.01. Nodes, representing the gene CNVs, are colored according to the MCL clustering (R package version 1.0) (*64*) of the gene CNVs absolute correlation values. The clustering was performed using respectively 2 and 3 for the expansion and inflation MCL parameters. The centroid and the standard deviation were computed for each cluster. Nodes mapping more than 2 standard deviations away from the cluster centroid were labeled as outliers and not shown. The network was produced using the R library igraph (R package version 1.1.2) (*65*), using the “layout_with_fr” function, placing vertices on the plane using the force-directed layout algorithm by Fruchterman and Reingold (*66*).

#### Phyogenetic analysis

The phylogenetic tree in **Fig. 1E** and the cladogram in **Fig. S5** include respectively the 191 ISC core isolates and the full 204 ISC set (*12*) and were calculated with raxmlHPC-PTHREADS (version 8.2.8) (*67*) with options ’-m ASC_GTRGAMMA --asc-corr=lewis’, with 10 starting trees and 10,000 rapid bootstrap runs. The program was run providing single nucleotide variant (SNV) positions returned by freebayes (version v1.0.1-2-g0cb2697) (*68*), and further filtered to produce high-quality predictions. Freebayes was run with options “--no-indels --read-indel-limit 0 --no-mnps --no-complex --read-mismatch-limit 3 --read-snp-limit 3 --hwe-priors-off --binomial-obs- priors-off --allele-balance-priors-off --min-alternate-fraction 0.05 --min-base-quality 5 --min-mapping-quality 50 --min-alternate-count 2 --pooled-continuous”. Then the variants mapping to the maxi-circle DNA or presenting multiple alternate alleles were discarded. Additionally, all variants with an alternate allele frequency < 0.9 were removed. Next, SNV predictions where the mean MAPQ of the reads supporting the alternate or the reference allele was < 20 were also removed. We filtered SNVs with sequencing coverage above or below 4 median absolute deviations (MADs) from the median chromosome coverage. Eventually, SNVs mapping inside homopolymers were filtered as described (*41*). To ease the visualization, the branch representing the reference genome was removed from the tree and the cladogram. The phylogenetic tree and the associated gene CNV heatmap were displayed with the R library ggtree (version 2.0.1) (*69*). A similar raxmlHPC-PTHREADS/freebayes approach was adopted to determine the evolutionary relationship between the P20 clones (**Fig. 2D**). As compared to the analyses in **Fig. 1E** and **S5**, we increased the number of starting trees (10, 000), used 10,000 standard bootstrap iterations, loosened the stringency of the alternate allele frequency filtering (< 0.1), and displayed the tree with iTol (version 5) (*70*).

#### RNAseq analysis

RNA was prepared and RNA sequencing (RNAseq) performed using an Illumina Hiseq 2000 platform and TruSeq v3 kits as previously described (*10*) We assessed the statistical significance of quantitative gene expression differences between WT clones and spontaneous NIMA-like kinase null mutant clones (spo-KO) with DESeq2 (version 1.14.1) (*71*). Genes with an adjusted p-value < 0.01 and a log2 fold change above 0.5 or below -0.5 were deemed differentially expressed. Maxi-circle DNA genes were not considered in the analysis. The gene expression was quantified by HTSeq (version 0.6.1) (*72*) while the reads were mapped as previously described (*41*).

To investigate post-transcriptionally regulated transcripts (**Fig. 3C**), we considered all the genes excluding the ones with 0 RNAseq read counts and the ones encoded on the maxi-circle DNAs. All HTSeq gene read counts estimated for RNAseq (RNA) and whole genome sequencing (DNA) were incremented by 1 pseudocount. For RNA and DNA data, the mean read counts in spo-KO clones was divided by the mean read counts in WT clones (representing the ratio scores that are displayed respectively on the y- and the x-axes in **Fig. 3C**). Both the numerator and denominator were normalized by the average library size (defined as the total number of reads mapping on all genes). The double-ratio score was estimated as the ratio between the RNA and DNA ratios. Genes with double-ratio score < 1 and p-value < 0.01 and were considered "destabilized" (i.e. less RNA than expected considering the DNA amount). Genes with double-ratio score > 1 and p-value < 0.01 were considered "stabilized" (i.e. more RNA than expected considering the DNA amount).

#### Gene Ontology (GO) term enrichment analysis

Cytoscape (version 3.7.2) (*73*) was used in combination with *BiNGO* (version 3.0.3) (*74*) to detect biological process GO terms enriched amongst the genes with significant double-ratio scores. In **Fig. 3D** node color mapping is ranging from yellow to dark orange to represent increasing significance levels, or lower adjusted p-values. White nodes are not significant. The BiNGO analysis was run using the “go-basic.obo” ontology file (release 2019-10-07) (*75, 76*) available from http://geneontology.org/docs/download-ontology/, and as a reference the set of 8,880 genes reporting a mean of normalized counts taken over all samples > 0 (“baseMean” metric in the DESeq2 analysis).

#### PCR analyses

The presence of a 10.3kb genomic deletion on chromosome 36 (coordinates 36:574,041 – 584,350) comprising a conserved NIMA kinase homolog (LdBPK_361580.1) was monitored by PCR amplification of a diagnostic fragment of 2kb. Briefly, genomic DNA was extracted from amastigotes purified from hamster spleen (*77*) or logarithmic promastigotes in culture using the DNeasy Blood and Tissue kit (Qiagen) according to manufacturer’s instructions. DNA quality was assessed by monitoring the 230/260 and 280/260 ratios and concentration was determined by spectrophotometry measuring the absorbance at 260nm. Genomic DNA was subjected to PCR analysis using LongAmp kit (New England Biolabs) according to the manufacturer’s instructions. Primers were designed targeting genomic sequence from either side of the deleted area (forward primer (F): AGACAGACAGCAATGCGTGAT; reverse primer (R): ACAGACGTCTGTCCGTGCTT) or the NIMA kinase open reading frame (F: ATGTCGGGGGGTAACGCC; R: TCACTCCTTTGCGGGTAGAGCA). The PCR mix was composed of 1x LongAmp Taq reaction buffer, 300µM dNTP each, 0.4µM of each primer, 10ng of genomic DNA and 5 units of LongAmp Taq DNA polymerase. The thermal cycling conditions were: initial denaturation at 94°C for 30s, followed by 30 cycles of 94°C for 30s, 60°C for 30s and 65°C for 10min. A final extension step was performed for 10min at 65°C.

PCR was performed for the CRISPR/cas9-mediated gene deletion assay. Based on the sequence of the NIMA kinase target gene (LdBPK_361580.1), single-guide RNA (sgRNA) were amplified using Expand High Fidelity polymerase (Roche) with 0.2mM dNTP each, 1x Expand High Fidelity buffer with MgCl_2_ (15 mM final concentration), 2µM each of primers G00 (sgRNA scaffold) and either 5’-sgRNA primer (gaaattaatacgactcactataggTAGAAAGAACGAAGTGATCGgttttagagctagaaatagc) or 3’-sgRNA primer (gaaattaatacgactcactataggAAGAGGAAGAAGATCACCAAgttttagagctagaaatagc) in 20µl final volume. PCR steps were: 30s at 98°C followed by 35 cycles of 10s at 98°C, 30s at 60°C, and 15s at 72°C. Sterilization of PCR products was performed at 94°C for 5min. PCR amplification of the DNA donor was performed using 30ng of pT plasmid containing the blasticidin resistance cassette as template (*78*). The PCR mix contained 0.2mM dNTP each, 1x Expand high Fidelity buffer with MgCl_2_ (3.375mM final concentration), 3% DMSO, 1 unit of Expand High Fidelity polymerase and 2µM of each primer (F: CTCTCCACCTCTGAATCTACTACGGTTTTCgtataatgcagacctgctgc; R: CGCAAAGGGTGCCGCCACATCGCAAACGCGccaatttgagagacctgtgc) in 40µl final volume. PCR steps were: 5min at 94°C followed by 40 cycles of 30s at 94°C, 30s at 65°C, 2min 15s at 72°C followed by a final elongation step for 7min at 72°C. Sterilization of PCR products was performed at 94°C for 5 min.

For the heterozygote NIMA^+/-^ null mutant, validation was performed by PCR amplification of LdBPK_361580.1 open reading frame (NIMA_F and NIMA_R) using 10ng genomic DNA as template and the amplification of the blasticidin resistance cassette (F: atgcctttgtctcaagaagaatc and R: ttagccctcccacacataac) using the 11kb fragment surrounding the insertion locus (diagn_F and diagn_R) as template.

For transgenic expression assay, the NIMA kinase gene LdBPK_361580.1 was PCR amplified using the Expand high fidelity polymerase kit (Roche) and 30ng of genomic DNA of promastigotes as template with primers F: ACCCTCGAGATGTCGGGGGGTAAC and R: CGCCTTAAGTCACTCCTTTGCGGGTAGAG. The amplicon was first sub-cloned into pGEM-T (Promega), validated by sequencing, and inserted into pXNG (kindly given by Stephen Beverley, Washington University School of Medicine, St. Louis, MO, USA, (*79*)) to generate construct pXNG-NIMA following digestion by XhoI and AflIII (*80*). For the validation of the pXNG transfection in the heterozygote NIMA^+/-^ null mutant, PCR assay was performed targeting the nourseothricin resistance cassette (F: ACCGTCGACATGAAGATTTCGGTGAT; R: CGGTCTAGATTAGGCGTCATCCTGT) after extraction of circular DNA from promastigotes at logarithmic growth phase using the Nucleospin Plasmid kit (Macherey Nagel).

#### Null mutant analysis

Vector pTB007 was used, containing a T7 RNA polymerase gene, the humanized Streptococcus pyogenes Cas9 gene, and homology regions allowing for stable integration into the beta-tubulin locus (*78*). The plasmid was linearized with SbfI (NEB) and Hind-III (NEB) and dephosphorylated using the Antarctic phosphatase (NEB) for 30min at 37°C. The 12kb-fragment was extracted and purified from 0.8% agarose gel using the Wizard SV Gel and PCR clean-up system (Promega). Transfection of P2 promastigotes from logarithmic growth phase was performed using a BioRad Gene pulser (*81*). After centrifugation (1600*g*, 10min, room temperature), cells were resuspended in Cytomix (0.15mM Cacl_2_, 120mM KCl, 10mM KH_2_PO_4_, 5mM MgCl_2_, 25mM HEPES (pH 7.5), 2mM EDTA (pH 7.6)) and 5.10^7^ cells were electroporated by two pulses at 1,500V, 25µF, infinite resistance. The next day, hygromycin B (Invitrogen) was added to select for transgenic cells (30µg/ml final concentration). Virulence of the transgenic line (termed Ld1S PT007) was maintained by two successive passages in hamsters as described (*77*) and recovered parasites were grown in the presence of hygromycin B. To generate heterozygous NIMA*^+/-^* null mutants, Ld1S PT007 promastigotes from logarithmic growth phase were transfected with the Biorad Gene Pulser combining sterilized PCR constructs and sgRNA. After centrifugation (1600*g*, 10min, room temperature), cells were resuspended in Cytomix and 3x10^7^ cells were electroporated using the following conditions: 900V, 50µF, infinite resistance, two pulses. The next day, blasticidin S hydrochloride (Sigma) was added to select for resistant cells (20 µg/ml final concentration) and isolate heterozygous NIMA*^+/-^* null mutants.

To generate the addback control, NIMA*^+/-^* promastigotes from logarithmic growth phase were transfected in electroporation buffer containing 90mM sodium phosphate, 5mM potassium chloride, 0.15mM calcium chloride, 50mM HEPES, pH 7.3 (*82*) using the program X-001 of the Amaxa Nucleofector IIb (Lonza) with either 5µg pXNG Mock or pXNG-NIMA DNA. For selection of transfected cells, nourseothricin (Sigma) was added 24 hours after electroporation at 100µg/ml final concentration.

#### Methylation (Nm) and pseudouridine (Ψ) rRNA modification analyses

We quantitatively mapped the position of individual Nm sites in *L. donovani* rRNA based on Cryogenic electron microscopy (Cryo-EM) (*40*) and mass-spectrometry (MS) (*83*) (**Table S20**). The snoRNAs predicted to guide Nm site were identified by homology to *L. major* snoRNAs (*84*) (**Table S20**). The relative methylation score (RMS score) of each individual Nm site was measured for both cell culture P2 and P135 (**Table S20**). Three biological replicates of P2 and two biological replicates of P135 were used for the analysis.

The fold change of Ψ−ratio between the reads obtained from Ψ-seq (*19*) at specific modification sites after and before 1-Cyclohexyl-3-(2-morpholinoethyl)carbodiimide metho-p-toluenesulfonate, 95%, ACROS Organics™ (CMC) (Fisher Scientific Acros 111360050) treatment of P135/P2 (**Formula 1**) was calculated across the rRNA based on four independent replicates. Only sites with log2 fold change >1.3 in all replicates were considered as hypermodified.

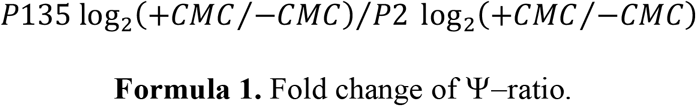

---

**Fig. S1.**
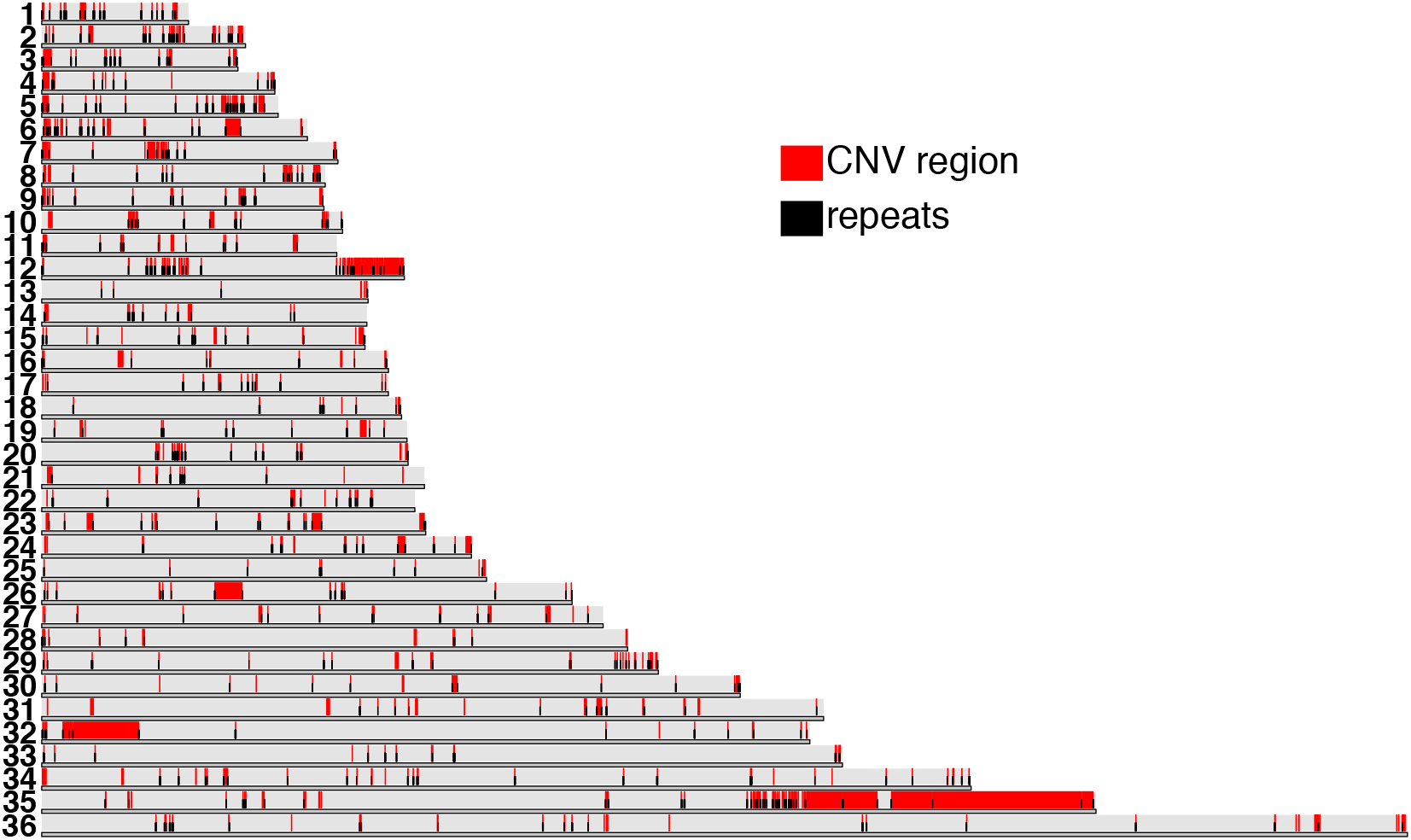
Genomic distribution of CNV regions and DNA sequence repeats. *Leishmania* chromosomes of the LdBPKv2 (*7*) are represented as stacked bars with karyoploteR (R package version 1.6.3) (*85*). The collapsed CNV sequence bins are shown in red. The repeat elements significantly associated with the CNV regions and mapping within their boundaries (1kb toward the outside and 150bp toward the inside of each collapsed CNV region) are shown in black.

**Fig. S2.**
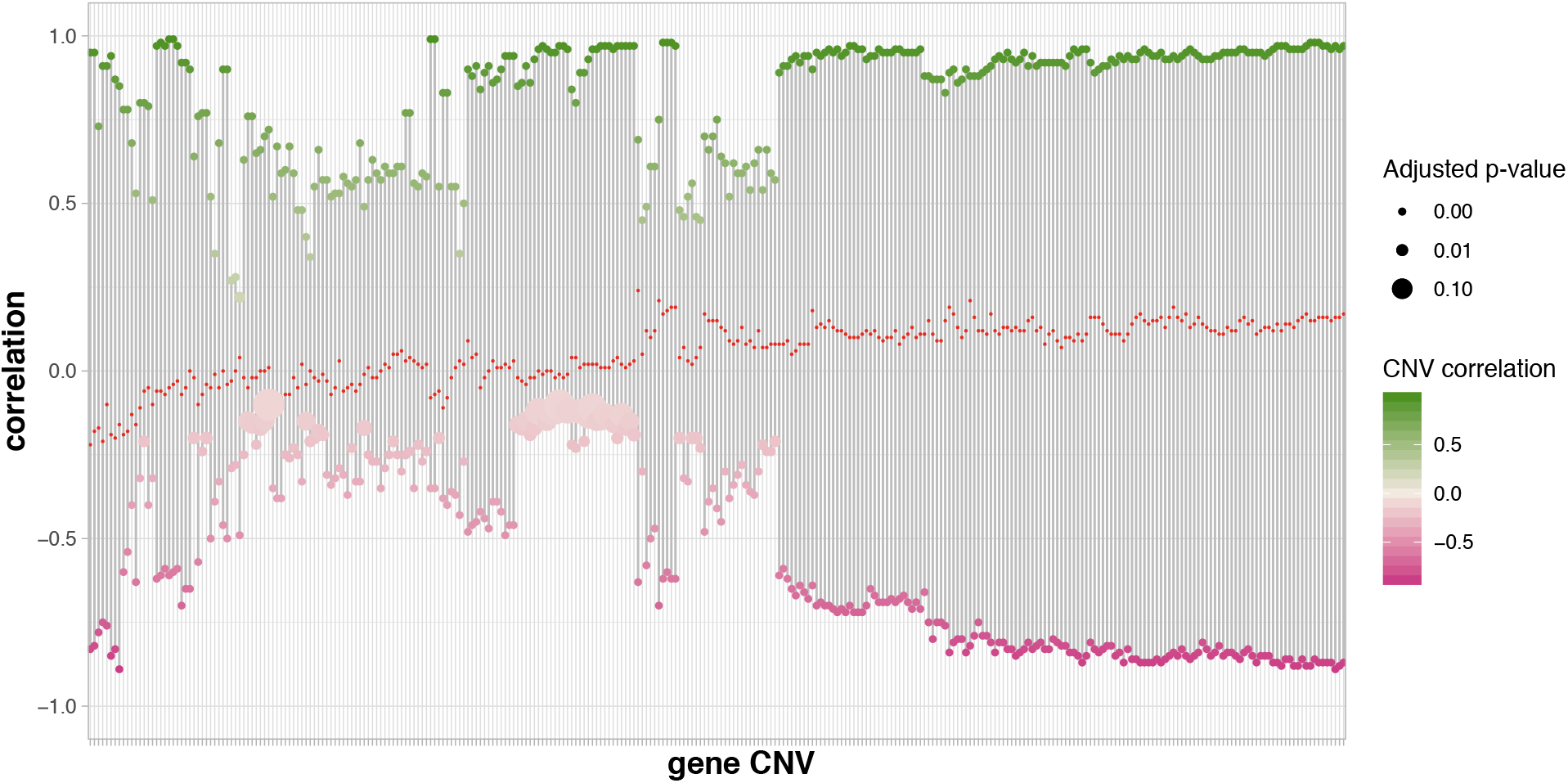
Distribution of correlation values measured between gene CNVs pairs. Each column indicates a different gene CNV. The *y*-axis reports the Pearson correlation value of the most positively (top, green) and most negatively (bottom, pink) correlated gene CNVs. The color intensity and the size of the dots match respectively to the correlation and the adjusted p-value scores. The red notches in the center indicate the median correlation scores.

**Fig. S3.**
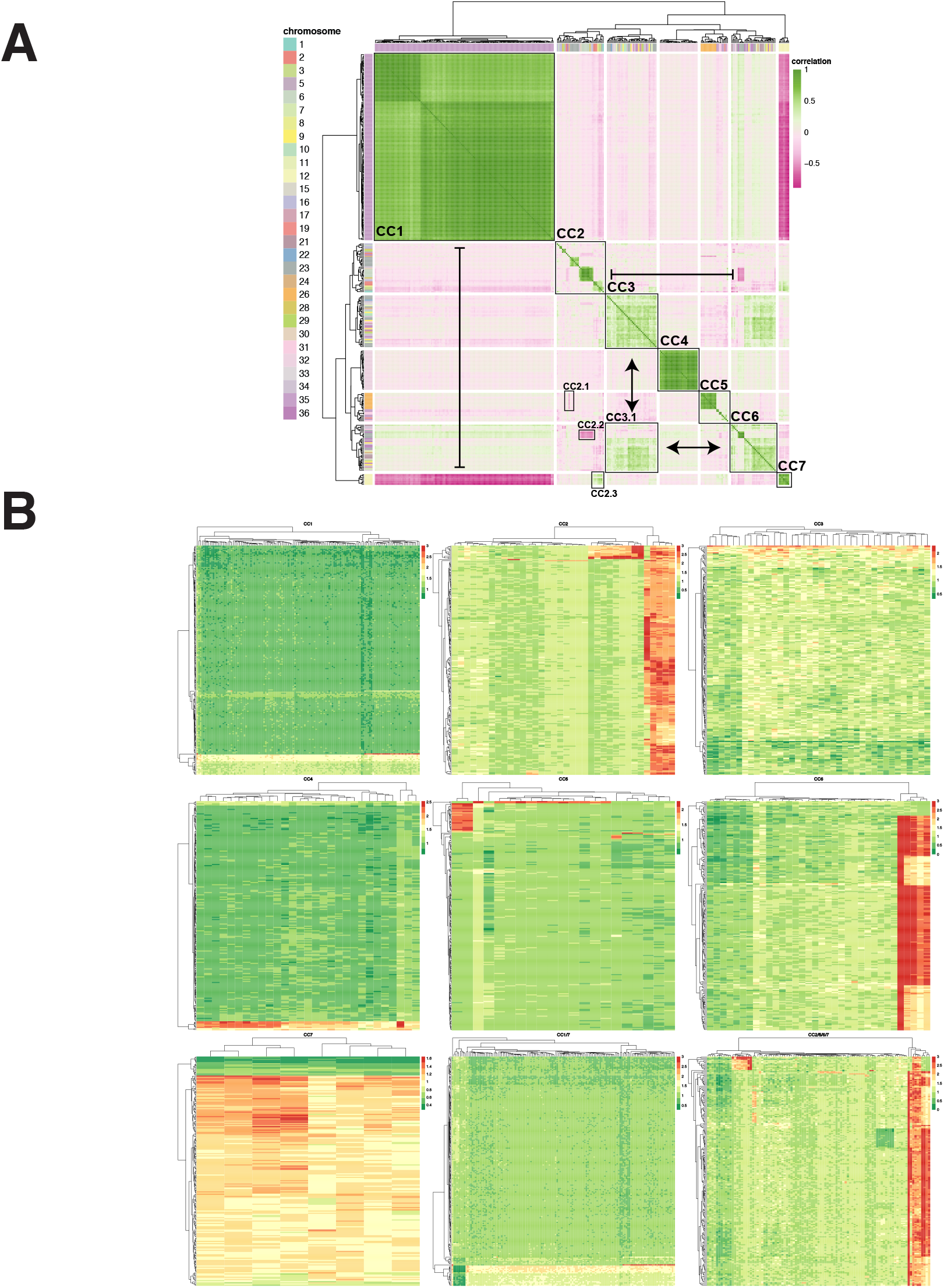
All-vs-all gene CNV correlation. (**A**) The heatmap shows gene CNVs on both axes. The color range reflects the Pearson normalized coverage correlation value of gene CNVs across 204 samples. The side ribbon indicates the chromosomal localization of the gene CNV as defined in the legend. Black boxes highlight correlations clusters (CC). The arrows indicate higher-level interactions between clusters. Pointy and flat arrowheads indicate respectively correlation and anti-correlation between clusters. (**B**) The heatmaps show the normalized coverage values of the gene CNVs (columns) belonging to the correlation clusters defined in panel A with respect to the 204 samples (rows). Green and red indicate respectively low and high coverage values as indicated in the legends. To ease the visualization, all scores of > 3 were assigned to a value of 3. The two last heatmaps show a combination of multiple CCs. In CC1/7 the columns include both CC1 and CC7 gene CNVs. In CC2/5/6/7 the columns include CC2, CC5, CC6, CC7 gene CNVs.

**Fig. S4.**
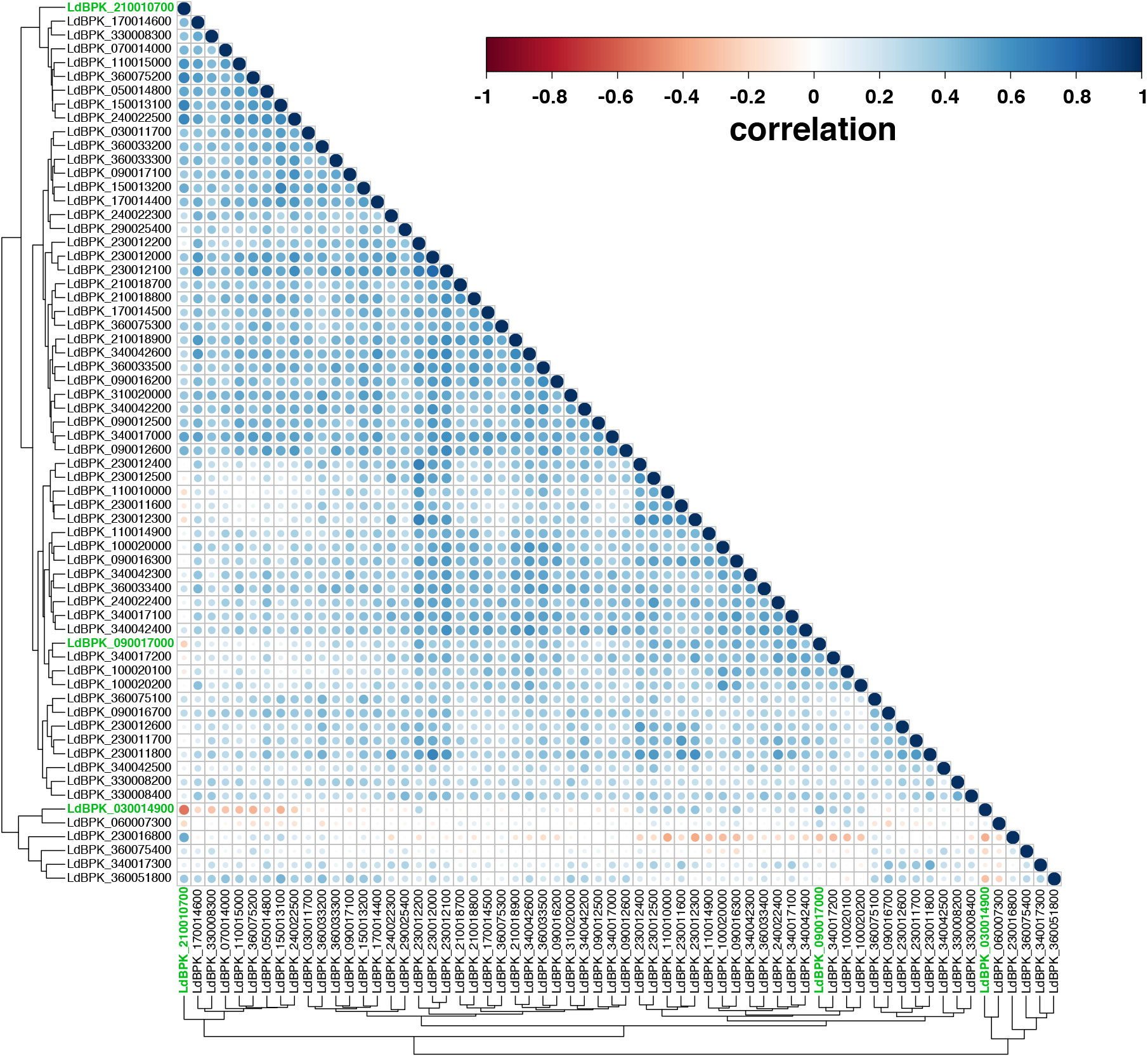
tRNA gene CNV correlation map. Both the *x-* and *y*-axes report gene CNVs. In green are labeled three genes that are network-hubs as judged by their high connectivity (LdBPK_210010700, rRNA; LdBPK_090017000, tRNA; LdBPK_030014900, eukaryotic translation initiation factor 2 subunit alpha). The color range of the dots reflects normalized coverage correlation value of the gene CNVs. The level and direction of correlation are indicated by both dot size and color (red, negative correlation; blue, positive correlation).

**Fig. S5.**
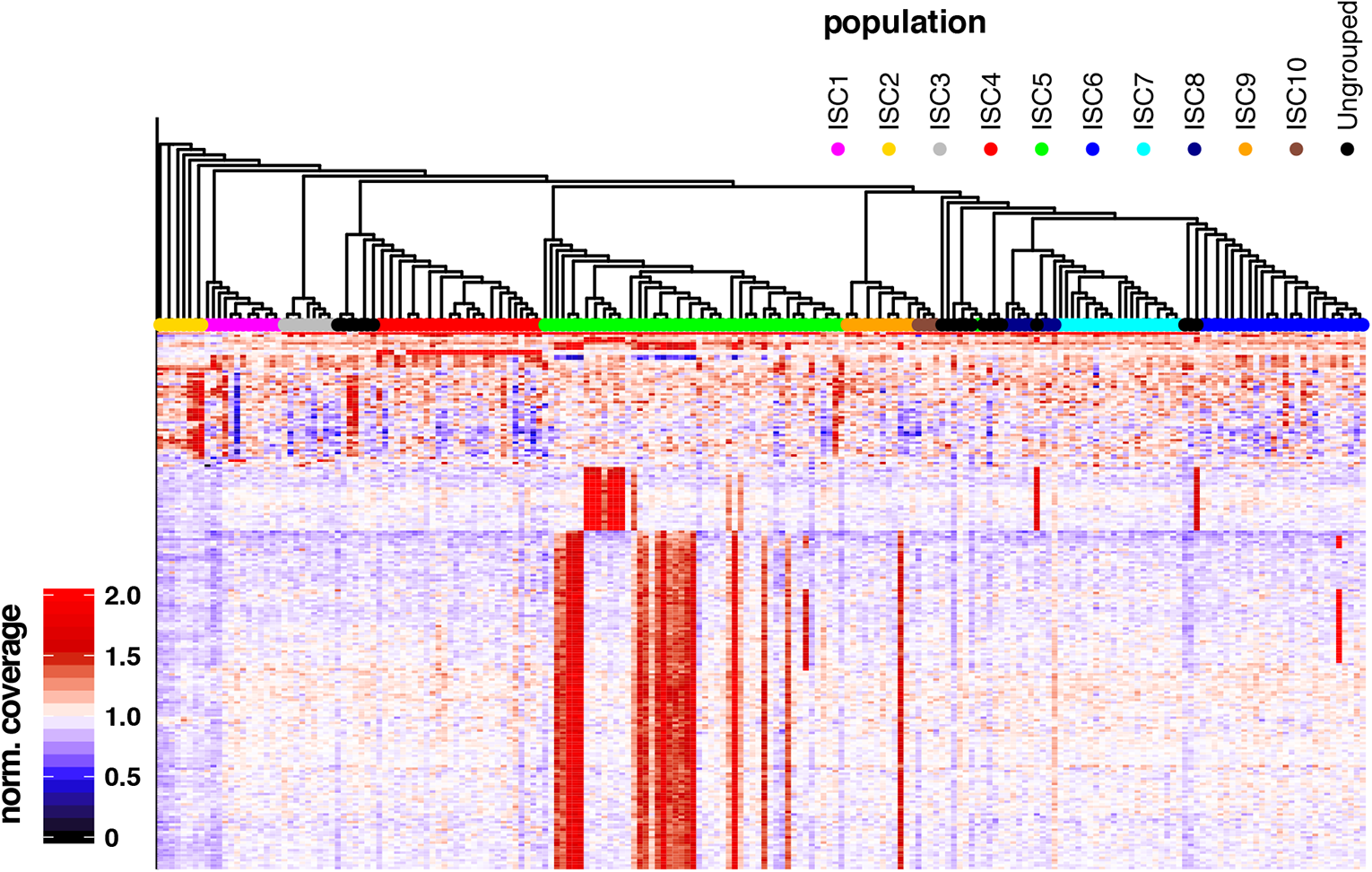
Polyphyletic distribution of gene CNVs. Cladogram based on SNVs (>90% frequency) (upper panel) and heatmap (lower panel) generated for the 204 ISC isolates. The cladogram topology demonstrates the relationships between isolates. The branch length does not reflect genetic distance. The heatmap has the same layout as in Fig. 1E.

**Fig. S6.**
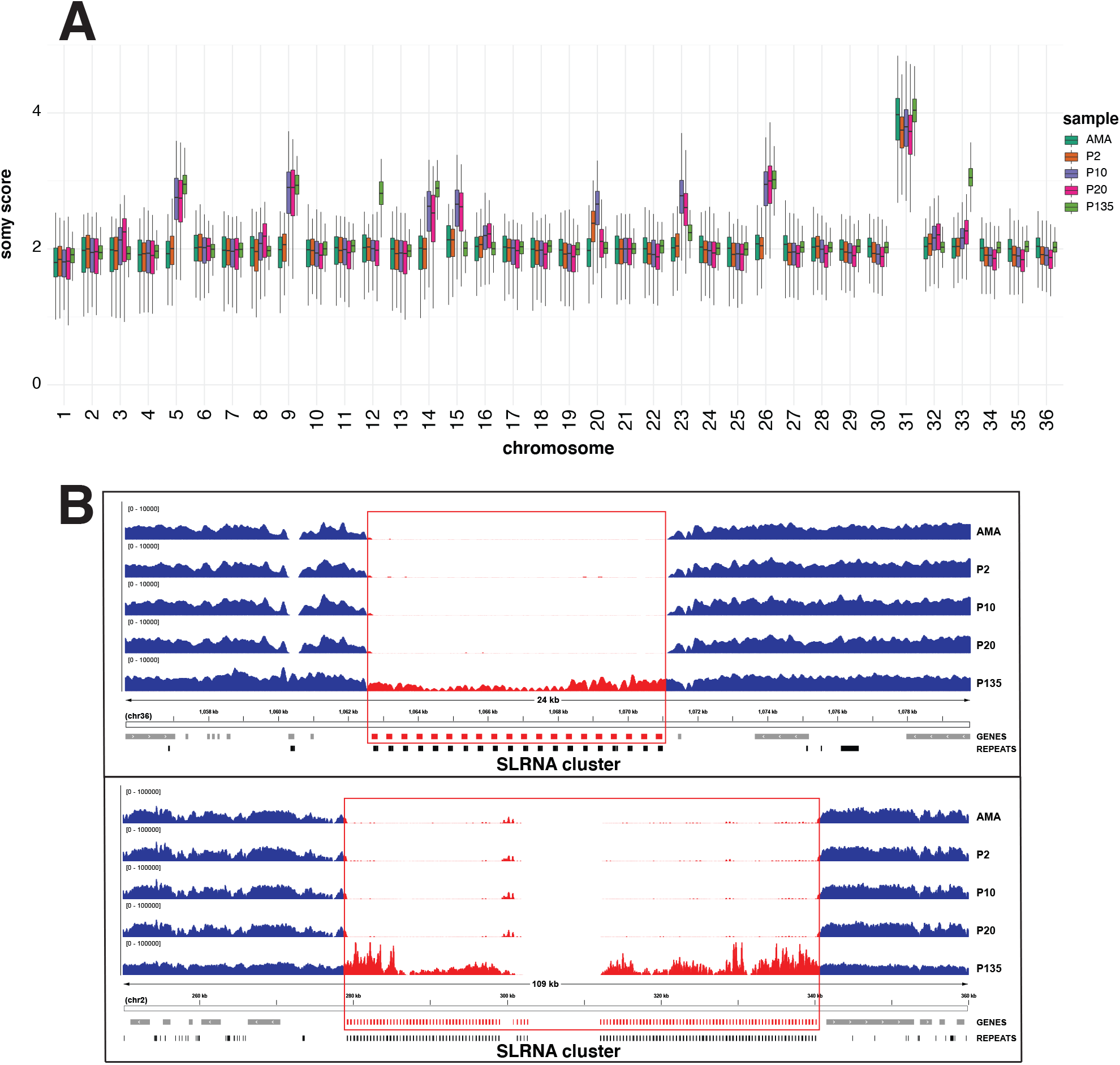
Chromosome and gene copy number comparison of splenic amastigotes (AMA) and derived promastigotes at different culture passages P2, 10, 20 and 135. (**A**) The box plots demonstrate the normalized sequencing coverage distributions of each chromosome. Bases where more than 50% of the reads have a MAPQ score lower than 50 were not considered. The somy score measured on the *y*-axis is an estimation of the chromosome copy number (see methods). Chromosome 21 showed a steady disomic level across this sample set and thus was used to normalize the read depth of the other chromosomes. (**B**) Examples of detected CNV gene clusters. Each panel illustrates a genome browser representation of the sequencing depth measured in the samples (see methods). Gene annotations and the predicted repetitive elements are indicated.

**Fig. S7.**
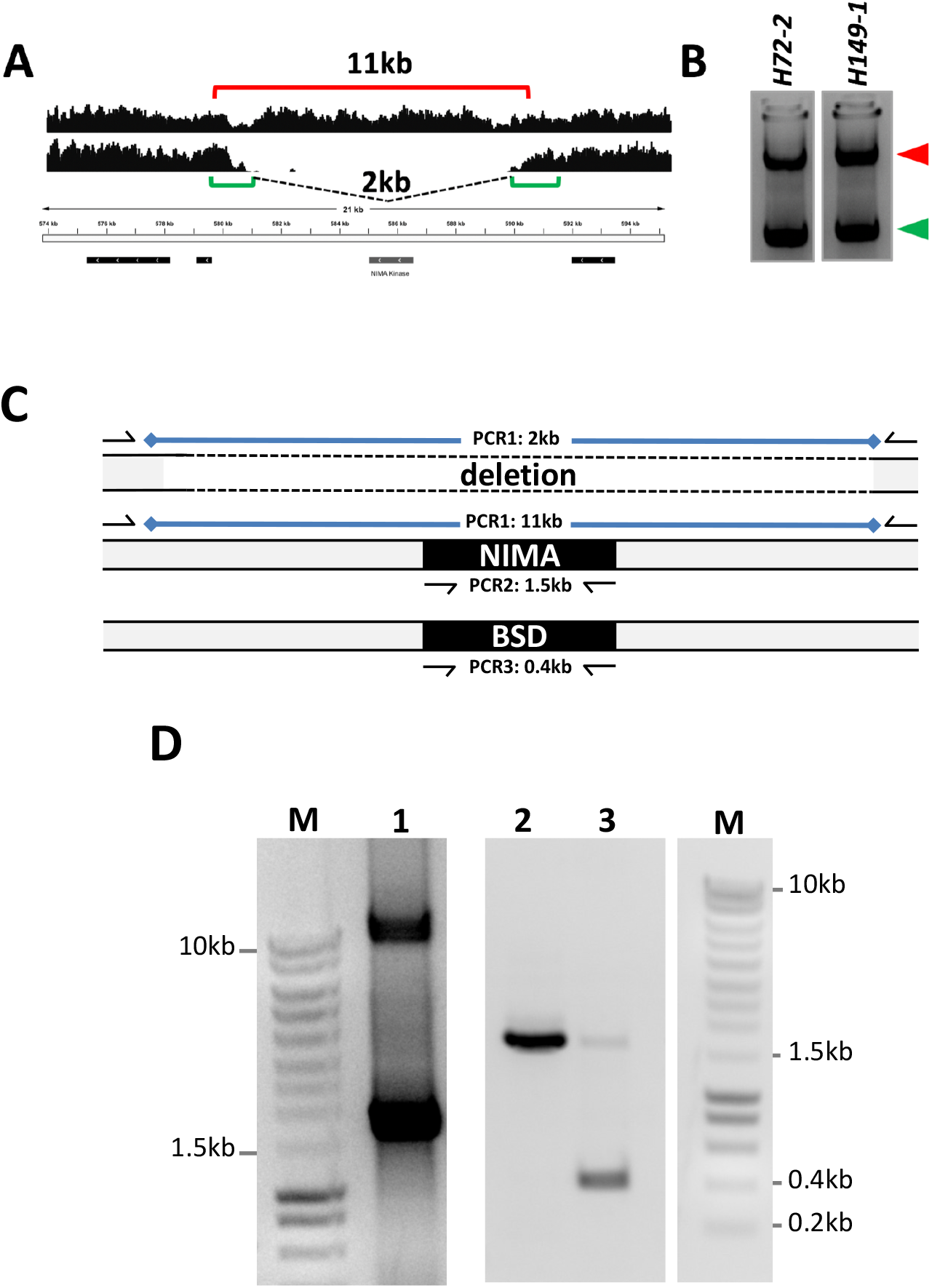
Generation of heterozygous NIMA null mutants using CRISPR/Cas9 gene editing. (**A**) Schematic representation of the NIMA locus, indicating the 11kb and 2kb PCR fragments diagnostic for the WT and deleted locus, respectively. (**B**) Gel electrophoretic separation of PCR fragments obtained from two independent splenic isolates purified from hamsters. The arrows heads indicate the 11kb (red) and 2kb (green) PCR fragments. (**C**) Schematic representation of the PCR amplification strategy. Given the size of the genomic fragment under investigation and the repetitive sequence surrounding the deletion site, a nested PCR protocol was applied first amplifying the target locus (PCR1) before reamplification of the open reading frames for the NIMA-related kinase gene (NIMA, PCR2) or the blasticidin resistance gene (BSD, PCR3). The PCR primers are indicated by the arrows. (**D**) Gel electrophoretic separation of PCR fragments obtained from a culture-adapted parasite population at *in vitro* passage 3 before gene editing using primer pair for PCR1 (lane 1), and of a representative NIMA+/-heterozygous clone using primer pairs PCR2 and PCR3 (lanes 2 and 3). The molecular weights of standard DNA marker fragments (M) are indicated.

**Fig. S8.**
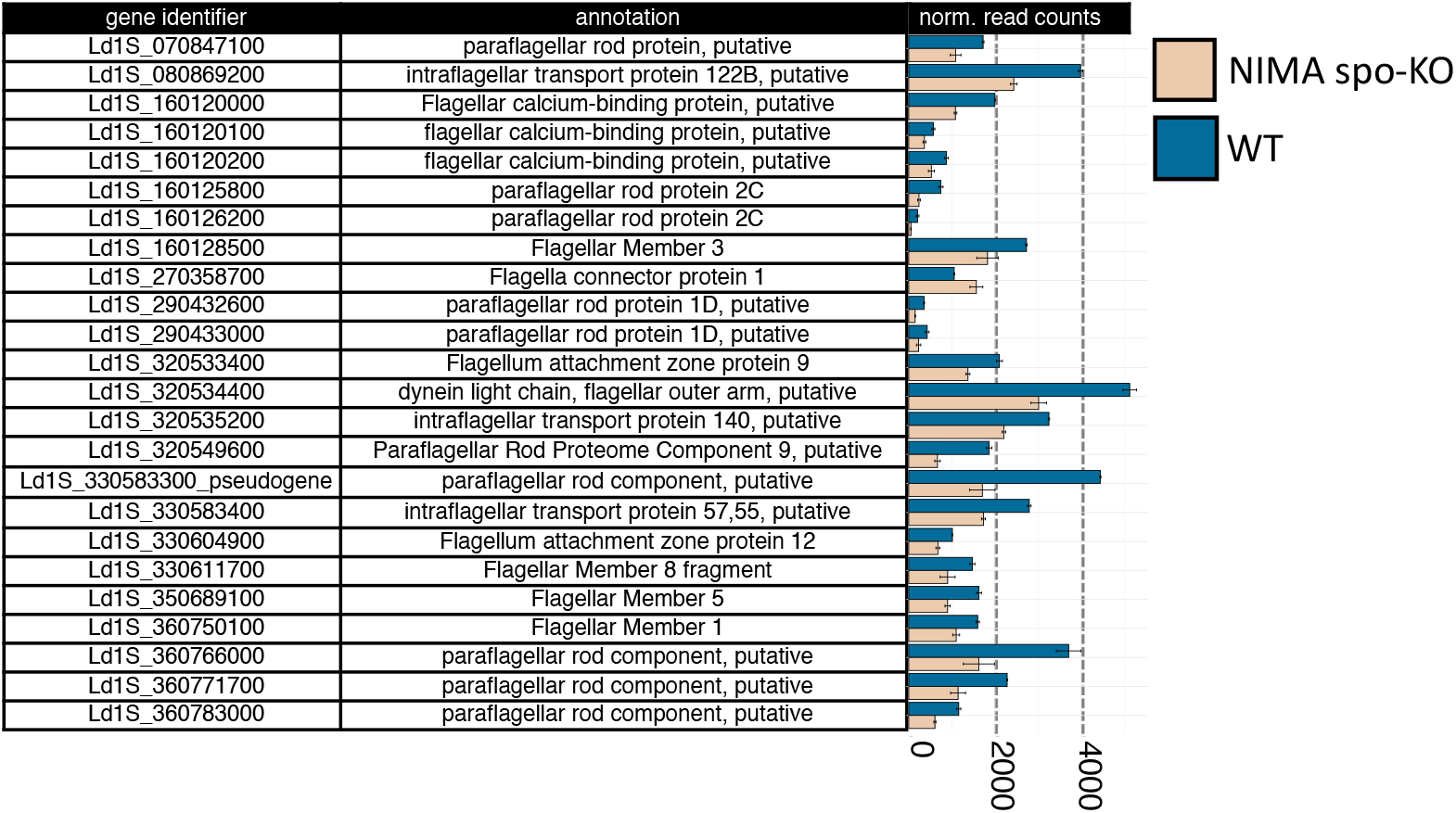
Functional analysis of genes differentially expressed between spo-KO and WT clones. Bar plot (right panel) showing the normalized mean RNAseq read counts of the genes related to flagellar function that were differentially expressed between WT (mean of two clones) and the six NIMA-kinase spo-KO clones. For each entry, the gene identifier and the annotated function are shown (left panel). The error bars indicate the standard deviations from the mean.

**Fig. S9.**
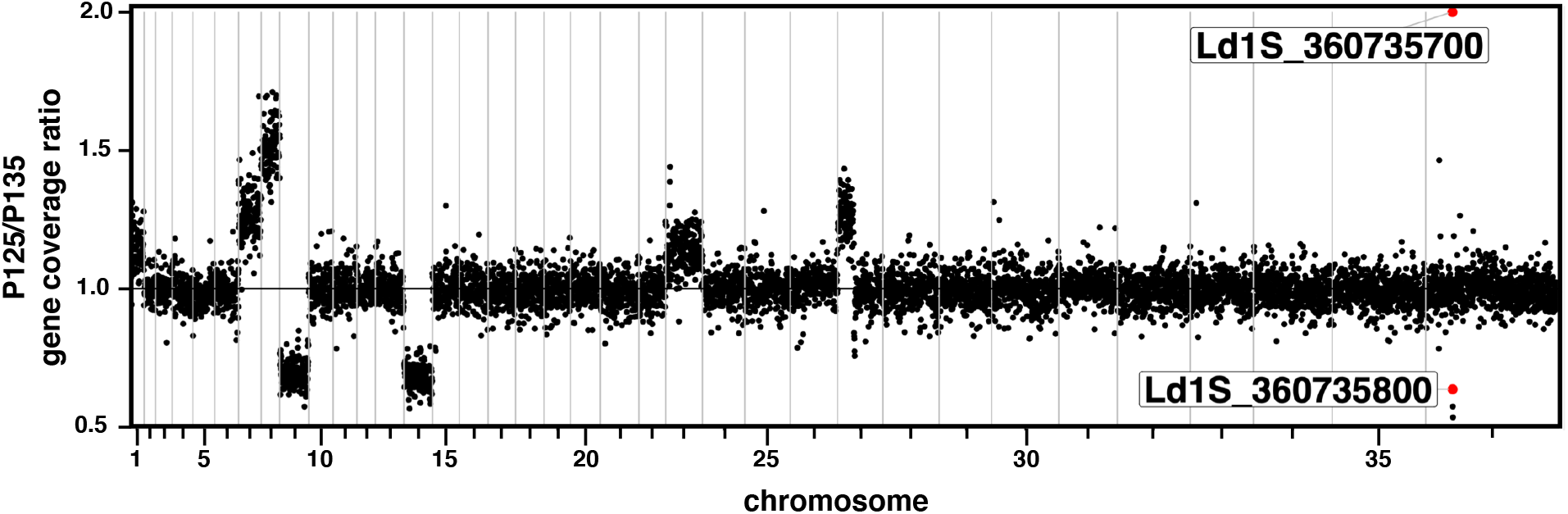
Different parasite populations show distinct evolutionary trajectories. (**A**) Genome-wide gene coverage ratio between sample P125 and P135. Each dot represents a different gene. Vertical lines mark chromosome boundaries. The deleted NIMA-kinase gene (Ld1S_360735700) and its homologue (Ld1S_360735800) are indicated by the red dot. To ease plot readability, all scores of > 2 were assigned to a value of 2.

**Fig. S10.**
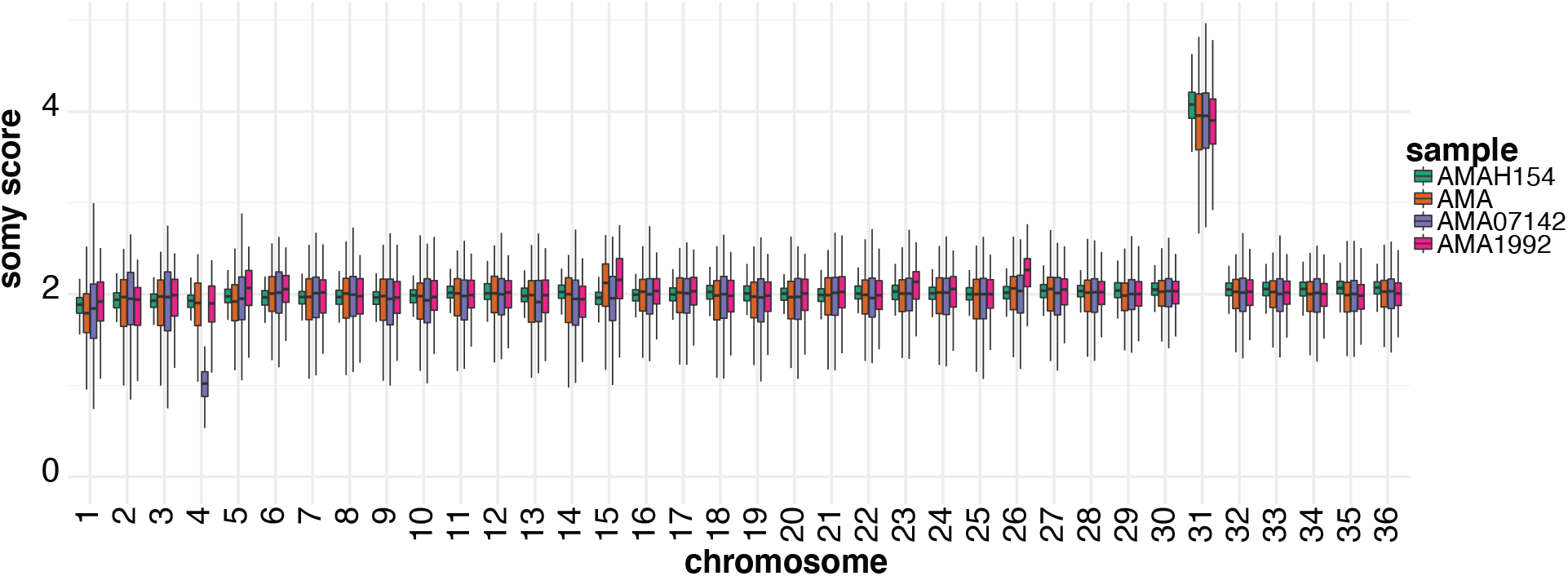
Chromosome ploidy analysis. The plot shows the karyotypic heterogeneity of four independent amastigote isolates (AMAH154, AMA, AMA07142, AMA1992). Same layout as in **Fig. S6A**, but the stable disomic chromosomes used for normalization was 25.

**Fig. S11.**
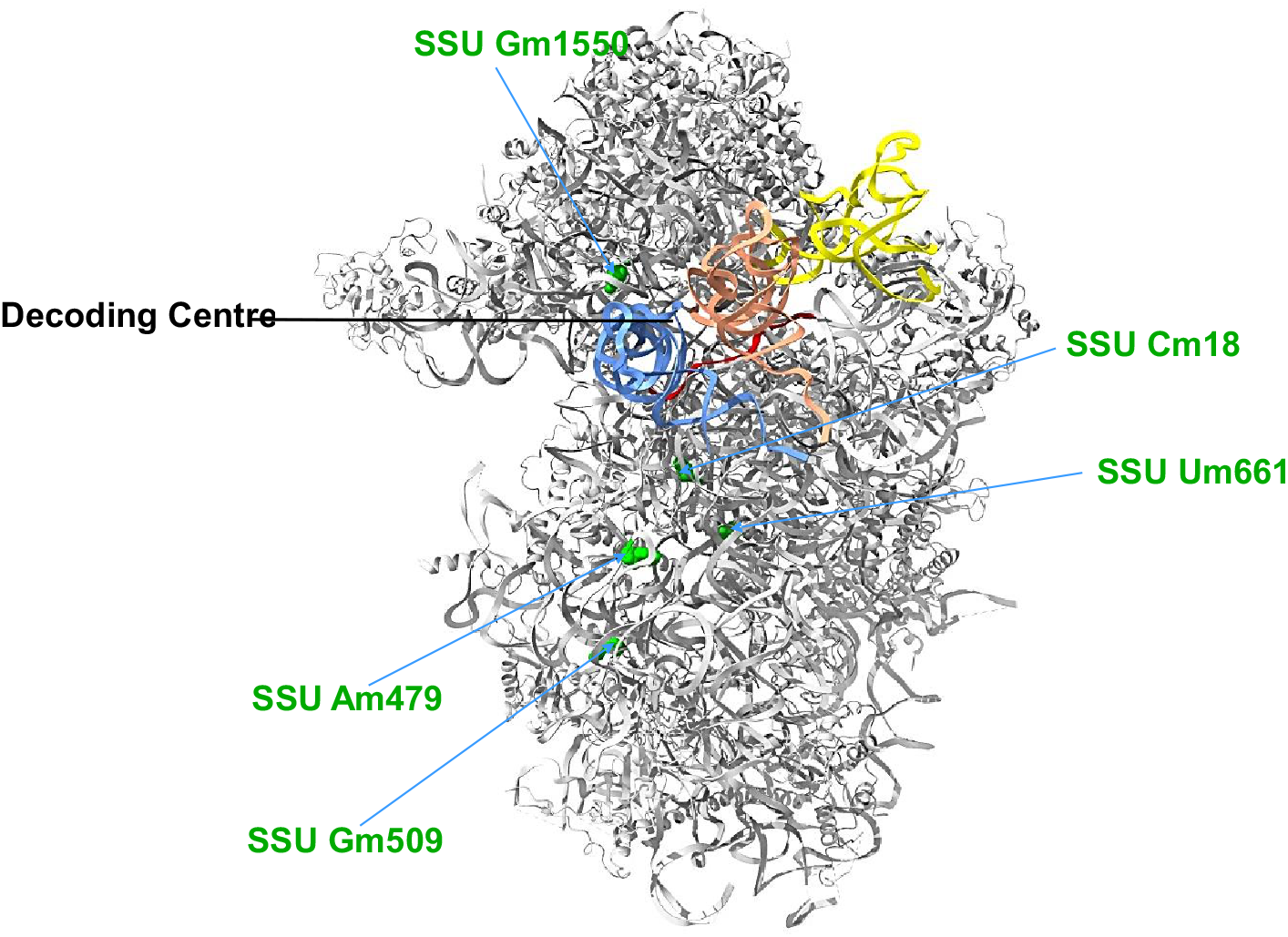
3D-structure of the ribosome small subunit (SSU) depicting the hypermodified Nm sites. The hypermodified Nm sites are shown in green. The location of the decoding center is indicated. The tRNAs positioned in the ribosome A (blue), P (orange) and E (yellow) sites are shown. The mRNA being translated in the ribosome is indicated in red. The representation is based on the previously deposited *L. donovani* ribosome SSU cryo-EM coordinates (Protein Data Bank (*39*) accession 6AZ1) (*40*).

**Data S1. (separate file)**

**Table S1: Mapping statistics of the dataset for the 204 clinical isolates.**

**Data S2. (separate file)**

**Table S2: Position and normalized coverage values of the collapsed CNV regions.**

*Footnote: The values reported for each sample indicate the normalized collapsed CNV coverage*.

**Data S3. (separate file)**

**Table S3: Repetitive elements.**

*Footnotes: The columns "GC%" and "N%" indicate respectively GC (guanine-cytosine) and the undetermined bases content. The "longest ORF" column indicates the length of the amino acid sequence of the longest open reading frame computed with EMBOSS getorf (version 6.6.0)* (*86*) *with option “-find 1”. (*) The observed count. (**) The expected count based on the sampled CNV boundaries. (***) The value at the 5% percentile of random samples. (****) The value at the 95% percentile of random samples. (*****) The standard deviation of random samples. (******) The fold enrichment, given by the ratio observed / expected. (*******) Log2 of the fold enrichment value. (********) The p-value of enrichment/depletion. (*********) The multiple-testing corrected p-value*.

**Data S4. (separate file)**

**Table S4: Most positive and most negative LdBPK gene coverage correlations.**

**Data S5. (separate file)**

**Table S5: Correlation map data.**

*Footnote: The first three columns and the first three rows indicate respectively LdBPKv1 gene identifiers, gene annotations and LdBPKv2 gene identifiers*.

**Data S6. (separate file)**

**Table S6: Correlation network data.**

*Footnote: The first three columns and the first three rows indicate respectively LdBPKv1 gene identifiers, gene annotations and LdBPKv2 gene identifiers*.

**Data S7. (separate file)**

**Table S7: Network hub genes.**

**Data S8. (separate file)**

**Table S8: Collapsed CNVs showing highest read depth in the 204 L. donovani field isolates.**

*Footnote: (*) Maximum measured coverage of the top 30 collapsed CNV detected in genic areas*.

Data S9. (separate file)

**Table S9: Genes amplified in phylogenetically distinct clades.**

*Footnote: The values reported for each sample indicate the normalized gene coverage*.

**Data S10. (separate file)**

**Table S10: Normalized gene coverage values across ISC samples.**

*Footnote: The values reported for each sample indicate the normalized gene coverage. (*) Difference between maximum and minimum normalized gene coverage*.

**Data S11. (separate file)**

**Table S11: Polysomy level analyses of two sets of Ld1S samples: (i) AMA to P135 isolates;**

**(ii) four AMA isolates.**

**Data S12. (separate file)**

**Table S12: Ld1S annotation.**

*Footnote: (*) Gene Ontology (GO) identifiers of the biological process (BP), molecular function (MF) and cellular component (CC). (**) LdBPKv1 homologs predicted with OrthoFinder. Genes with no significant match were searched with NCBI-blastn. One-to-many homologs are reported with comma separated gene identifiers*.

**Data S13. (separate file)**

**Table S13: Definitions of gene clusters in two sets of Ld1S samples: (i) AMA-P135 isolates;**

**(ii) P20 clones isolates.**

**Data S14. (separate file)**

**Table S14: Gene CNVs and cluster CNVs detected during Ld1S culture adaptation.**

*Footnotes: (*) The AMA, P2, P10, P20 and P135 columns indicate the normalized gene coverage in the respective samples. (**) “increasing” gene CNVs have a P135 normalized coverage increase of at least 0.5 with respect to AMA. “decreasing” gene CNVs have a P135 normalized coverage decrease of at least 0.5 with respect to AMA. All the other gene CNVs are labeled as “transient”. (***) The number of genes in the gene cluster (clstr). Individual (non-cluster) genes have by default a value of 1. (****) Gene Ontology (GO) identifiers of the biological process (BP), molecular function (MF) and cellular component (CC). (*****) The genes belonging to each cluster CNV are reported as separate entries sharing the same attributes.*

**Data S15. (separate file)**

**Table S15: Gene CNVs and cluster CNVs detected in the eight Ld1S P20 clones.**

*Footnotes: (*) The CL1, CL3, CL4, CL6, CL7, CL8, CL9 and CL10 columns indicate the normalized gene coverage in the respective samples. (**) The number of genes in the gene cluster (clstr). Individual (non-cluster) genes have by default a value of 1. (***) Gene Ontology (GO) identifiers of the biological process (BP), molecular function (MF) and cellular component (CC). (****) The genes belonging to each cluster CNV are reported as separate entries sharing the same attributes*.

**Data S16. (separate file)**

**Table S16: Single nucleotide variants (SNVs) in Ld1S P20 clones.**

*Footnotes: (*) Number of alternate observations, (**) Total read depth at the locus, (***) Alternate allele quality sum in phred, (****) Reference allele quality sum in phred, (*****) Mean mapping quality of observed alternate alleles, (******) Mean mapping quality of observed reference alleles, (*******) Reference base and alternate allele, (********) context sequence of the variant (+/-5bp), (*********) SnpEff predicted variant effect*.

**Data S17. (separate file)**

**Table S17: Data of the differential expression analysis.**

*Footnotes: DESeq2 was utilized to test differential gene expression between the NIMA spo-KO and the WT clones groups. Significant genes (adjusted p-value < 0.01) with a log2 fold change >*

*0.5 were considered “enriched” in the NIMA spo-KO group, while those with a log2 fold change*

*< -0.5 were considered “depleted” in the NIMA spo-KO group. Maxi-circle genes were not considered in the analysis. (*) Mean of normalized counts, taken over all samples, (**) log2 fold change between the groups, (***) standard error of the log2 fold change estimate, (****) Wald statistic, (*****) Wald test p-value, (******) Benjamini-Hochberg adjusted p-value*.

**Data S18. (separate file)**

**Table S18: DNA and RNA double-ratio data.**

*Footnotes: (*) Whole genome sequencing (DNA) ratio: the mean read counts in spo-KO clones is divided by the mean DNA read counts in WT clones. Both the numerator and denominator are normalized by the average library size. (**) RNAseq (RNA) ratio: the mean read counts in spo-KO clones is divided by the mean RNA read counts in WT clones. Both the numerator and denominator are normalized by the average library size. (***) Double-ratio score: the ratio between the RNA ratio and the DNA ratio. (****) P-values of the double-ratio scores. (*****)*

*Genes with double-ratio score p-value < 0.01 are considered "destabilized" or "stabilized" for double-ratio scores < or > 1, respectively. (******) Gene Ontology (GO) identifiers of the biological process (BP), molecular function (MF) and cellular component (CC)*.

**Data S19. (separate file)**

**Table S19: P125/P135 gene coverage ratio values.**

*Footnote: (*) The P125 and P135 columns indicate the gene coverage normalized by the median gene coverage in the respective samples*.

**Data S20. (separate file)**

**Table S20: The complete stoichiometry of Nm sites in L. donovani rRNA.**

*Footnote: Data are shown as mean ± SEM (standard error of mean). The fold change (FC) of RMS of individual Nms is presented and the hypermodified sites showing FC > 2 are highlighted. Note that we have not identified all the snoRNAs guiding the existing Nm modification. (*) The rRNA unit (LSU, SSU or 5.8S) and the nucleotide carrying the Nm modification are reported*.

**Data S21. (separate file)**

**Table S21: The relative fold-change of Ψs in L. donovani rRNA between P135 and P2 parasites.**

*Footnote: (*) Four independent replicates (R). Only sites with log2 fold change (FC) >1.3 in all replicates are considered as hypermodified sites and are highlighted in pink*.

## Notes

### Competing Interest Statement

The authors have declared no competing interest.

