## Supplementary material for "Genome instability drives epistatic adaptation in the human pathogen *Leishmania*": Table S20

**Table S20.** **The complete stoichiometry of Nm sites in *L. donovani* rRNA.**

| **Nm position (*)** | **snoRNA** | **Nm** | **Ref** | **P2-RMS** | **P135-RMS** | **FC P135/P2** |
| --- | --- | --- | --- | --- | --- | --- |
| 5.8S_162 | LD35Cs1C2 | Am | Cryo-EM | 0.88+/-0.01 | 0.88+/-0.01 | 1.00 |
| 5.8S_43 | LD26Cs2C1 | Am | Cryo-EM | 0.83+/-0.02 | 0.72+/-0.03 | 0.87 |
| 5.8S_7 | LD36Cs1C1 | Um | Cryo-EM | 0.81+/-0.07 | 0.62+/-0.05 | 0.77 |
| 5.8S_75 | LD23Cs1C2 | Gm | Cryo-EM | 0.55+/-0.08 | 0.24+/-0.00 | 0.43 |
| LSUa_1071 | LD33Cs1C2 | Um | MS | 0.86+/-0.01 | 0.68+/-0.06 | 0.80 |
| LSUa_1107 | LD33Cs1C2 | Um | Cryo-EM | 0.66+/-0.13 | 0.80+/-0.03 | 1.22 |
| LSUa_1190 | LD35Cs3C7 | Gm | Cryo-EM | 0.27+/-0.27 | 0.61+/-0.02 | 2.21 |
| LSUa_1253 | LD36Cs4C1A | Um | Cryo-EM | 0.55+/-0.15 | 0.76+/-0.04 | 1.39 |
| LSUa_1371 | LD36Cs2C3 | Um | Cryo-EM | 0.47+/-0.07 | 0.44+/-0.00 | 0.94 |
| LSUa_1526 | - | Gm | Cryo-EM | 0.83+/-0.07 | 0.45+/-0.02 | 0.55 |
| LSUa_1529 | - | Cm | Cryo-EM | 0.71+/-0.15 | 0.36+/-0.00 | 0.51 |
| LSUa_1541 | - | Am | Cryo-EM | 0.22+/-0.22 | 0.58+/-0.00 | 2.65 |
| LSUa_1542 | - | Gm | Cryo-EM | 0.73+/-0.17 | 0.32+/-0.07 | 0.44 |
| LSUa_1628 | - | Gm | Cryo-EM | 0.59+/-0.10 | 0.66+/-0.01 | 1.12 |
| LSUa_1661 | - | Um | Cryo-EM | 0.27+/-0.26 | 0.67+/-0.01 | 2.53 |
| LSUa_235 | LD33Cs1C4 | Am | Cryo-EM | 0.78+/-0.10 | 0.44+/-0.02 | 0.56 |
| LSUa_437 | - | Am | Cryo-EM | 0.15+/-0.03 | 0.22+/-0.02 | 1.52 |
| LSUa_48 | - | Um | Cryo-EM | 0.17+/-0.16 | 0.35+/-0.08 | 2.08 |
| LSUa_678 | LD35Cs3C3 | Am | Cryo-EM | 0.32+/-0.30 | 0.85+/-0.00 | 2.70 |
| LSUa_681 | LD33Cs2C2 | Am | Cryo-EM | 0.79+/-0.07 | 0.87+/-0.01 | 1.09 |
| LSUa_69 | - | Am | Cryo-EM | 0.33+/-0.03 | 0.15+/-0.05 | 0.43 |
| LSUa_695 | LD35Cs3C3 | Cm | Cryo-EM | 0.26+/-0.26 | 0.65+/-0.02 | 2.51 |
| LSUa_697 | - | Am | Cryo-EM | 0.26+/-0.13 | 0.13+/-0.11 | 0.48 |
| LSUa_845 | LD35Cs3C4 | Um | Cryo-EM | 0.67+/-0.08 | 0.75+/-0.00 | 1.12 |
| LSUa_847 | LD27Cs1C2 | Um | Cryo-EM | 0.42+/-0.21 | 0.44+/-0.12 | 1.06 |
| LSUa_856 | LD35Cs3C4 | Gm | Cryo-EM | 0.53+/-0.12 | 0.73+/-0.01 | 1.38 |
| LSUa_858 | LD27Cs1C2 | Am | Cryo-EM | 0.25+/-0.22 | 0.82+/-0.04 | 3.27 |
| LSUa_927 | LD14Cs1C2 | Am | Cryo-EM | 0.61+/-0.13 | 0.24+/-0.02 | 0.39 |
| LSUa_955 | LD25Cs1C3 | Am | Cryo-EM | 0.27+/-0.27 | 0.70+/-0.01 | 2.55 |
| LSUa_959 | LD34Cs1C1 | Gm | Cryo-EM | 0.78+/-0.14 | 0.28+/-0.05 | 0.35 |
| LSUb_1047 | - | Gm | Cryo-EM | 0.59+/-0.30 | 0.00+/-0.00 | 0.00 |
| LSUb_1068 | - | Am | Cryo-EM | 0.41+/-0.21 | 0.00+/-0.00 | 0.00 |
| LSUb_1078 | - | Um | Cryo-EM | 0.28+/-0.26 | 0.50+/-0.03 | 1.79 |
| LSUb_1079 | - | Gm | Cryo-EM | 0.37+/-0.14 | 0.36+/-0.00 | 0.96 |
| LSUb_1153 | - | Um | Cryo-EM | 0.12+/-0.06 | 0.19+/-0.07 | 1.51 |
| LSUb_1160 | - | Cm | Cryo-EM | 0.37+/-0.07 | 0.36+/-0.04 | 0.98 |
| LSUb_1186 | LD31Cs1C1 | Am | Cryo-EM | 0.94+/-0.03 | 0.81+/-0.02 | 0.86 |
| LSUb_1230 | LD25Cs1C1 | Gm | Cryo-EM | 0.85+/-0.05 | 0.63+/-0.03 | 0.74 |
| LSUb_1232 | - | Gm | Cryo-EM | 0.22+/-0.22 | 0.87+/-0.02 | 3.87 |
| LSUb_1249 | LD30Cs1C1 | Cm | Cryo-EM | 0.69+/-0.05 | 0.71+/-0.01 | 1.04 |
| LSUb_1254 | LD25Cs1C1 | Gm | Cryo-EM | 0.73+/-0.15 | 0.15+/-0.04 | 0.20 |
| LSUb_1318 | - | Cm | Cryo-EM | 0.20+/-0.11 | 0.36+/-0.01 | 1.81 |
| LSUb_1360 | LD33Cs1C3 | Um | Cryo-EM | 0.69+/-0.00 | 0.38+/-0.04 | 0.55 |
| LSUb_1361 | - | Gm | Cryo-EM | 0.40+/-0.03 | 0.20+/-0.02 | 0.49 |
| LSUb_1373 | LD27Cs1C1 | Am | Cryo-EM | 0.69+/-0.13 | 0.40+/-0.03 | 0.58 |
| LSUb_1385 | LD27Cs1C1 | Am | Cryo-EM | 0.86+/-0.02 | 0.76+/-0.01 | 0.88 |
| LSUb_1398 | LD35Cs3C6 | Cm | Cryo-EM | 0.28+/-0.28 | 0.72+/-0.02 | 2.53 |
| LSUb_1420 | LD35Cs2C1 | Um | Cryo-EM | 0.71+/-0.03 | 0.59+/-0.02 | 0.83 |
| LSUb_359 | LD35Cs1C1 | Cm | Cryo-EM | 0.66+/-0.07 | 0.46+/-0.01 | 0.69 |
| LSUb_382 | - | Am | Cryo-EM | 0.98+/-0.00 | 0.98+/-0.00 | 1.00 |
| LSUb_443 | LD18Cs1C2 | Cm | Cryo-EM | 0.50+/-0.09 | 0.56+/-0.01 | 1.12 |
| LSUb_502 | - | Am | Cryo-EM | 0.46+/-0.22 | 0.75+/-0.03 | 1.63 |
| LSUb_527 | LD26Cs1C1 | Am | Cryo-EM | 0.86+/-0.01 | 0.76+/-0.01 | 0.88 |
| LSUb_534 | LD36Cs1C2 | Gm | Cryo-EM | 0.94+/-0.02 | 0.85+/-0.02 | 0.91 |
| LSUb_560 | LD36Cs3C1 | Um | Cryo-EM | 0.69+/-0.12 | 0.85+/-0.02 | 1.22 |
| LSUb_570 | LD20Cs1C1 | Am | Cryo-EM | 0.66+/-0.19 | 0.00+/-0.00 | 0.00 |
| LSUb_572 | - | Am | Cryo-EM | 0.58+/-0.22 | 0.46+/-0.03 | 0.79 |
| LSUb_583 | LD36Cs1C3 | Cm | Cryo-EM | 0.22+/-0.13 | 0.45+/-0.02 | 2.07 |
| LSUb_591 | LD36Cs1C3 | Am | Cryo-EM | 0.45+/-0.01 | 0.42+/-0.01 | 0.93 |
| LSUb_604 | LD33Cs3C1 | Am | Cryo-EM | 0.35+/-0.27 | 0.79+/-0.02 | 2.25 |
| LSUb_628 | LD4Cs1C1 | Am | Cryo-EM | 0.82+/-0.03 | 0.63+/-0.03 | 0.77 |
| LSUb_641 | LD35Cs3C2 | Gm | Cryo-EM | 0.39+/-0.12 | 0.00+/-0.00 | 0.00 |
| LSUb_655 | LD27Cs1C3 | Gm | Cryo-EM | 0.64+/-0.06 | 0.60+/-0.02 | 0.93 |
| LSUb_665 | - | Am | Cryo-EM | 0.43+/-0.21 | 0.00+/-0.00 | 0.00 |
| LSUb_667 | LD35Cs3C2 | Um | Cryo-EM | 0.31+/-0.13 | 0.50+/-0.01 | 1.60 |
| LSUb_71 | LD26Cs1C3 | Gm | Cryo-EM | 0.80+/-0.10 | 0.35+/-0.01 | 0.44 |
| LSUb_710 | LD33Cs1C1 | Um | MS | 0.45+/-0.02 | 0.21+/-0.02 | 0.47 |
| LSUb_73 | - | Um | Cryo-EM | 0.14+/-0.14 | 0.72+/-0.02 | 5.16 |
| LSUb_95 | LD29Cs2C1 | Am | MS | 0.49+/-0.12 | 0.53+/-0.02 | 1.10 |
| SSU_115 | - | Cm | Cryo-EM | 0.56+/-0.04 | 0.39+/-0.05 | 0.70 |
| SSU_1464 | - | Gm | Cryo-EM | 0.10+/-0.03 | 0.13+/-0.00 | 1.33 |
| SSU_1478 | LD25Cs1C2 | Gm | Cryo-EM | 0.92+/-0.02 | 0.76+/-0.03 | 0.83 |
| SSU_1550 | LD23Cs1C1 | Gm | Cryo-EM | 0.31+/-0.31 | 0.84+/-0.03 | 2.70 |
| SSU_1599 | LD36Cs1C1 | Um | MS | 0.47+/-0.12 | 0.69+/-0.00 | 1.46 |
| SSU_1621 | LD33Cs1C4 | Um | Cryo-EM | 0.50+/-0.09 | 0.48+/-0.04 | 0.97 |
| SSU_1623 | LD5Cs1C1 | Gm | Cryo-EM | 0.55+/-0.19 | 0.52+/-0.04 | 0.95 |
| SSU_1647 | LD36Cs-1pC2 | Gm | Cryo-EM | 0.72+/-0.07 | 0.46+/-0.00 | 0.63 |
| SSU_1777 | LD14Cs1C1 | Um | Cryo-EM | 0.66+/-0.11 | 0.39+/-0.04 | 0.59 |
| SSU_18 | LD23Cs1C2 | Cm | Cryo-EM | 0.22+/-0.22 | 0.53+/-0.03 | 2.41 |
| SSU_1829 | LD35Cs3C5 | Gm | Cryo-EM | 0.85+/-0.09 | 0.34+/-0.03 | 0.41 |
| SSU_1833 | LD5Cs1C2 | Um | Cryo-EM | 0.48+/-0.17 | 0.65+/-0.04 | 1.36 |
| SSU_1865 | LD18Cs1C3 | Gm | Cryo-EM | 0.43+/-0.22 | 0.81+/-0.00 | 1.86 |
| SSU_1866 | - | Cm | Cryo-EM | 0.39+/-0.19 | 0.00+/-0.00 | 0.00 |
| SSU_1979 | LD18Cs1C3 | Um | Cryo-EM | 0.92+/-0.01 | 0.87+/-0.00 | 0.95 |
| SSU_2021 | LD5Cs1C4 | Am | Cryo-EM | 0.76+/-0.01 | 0.63+/-0.02 | 0.83 |
| SSU_2048 | LD18Cs1C1 | Um | Cryo-EM | 0.48+/-0.13 | 0.73+/-0.03 | 1.53 |
| SSU_2059 | LD18Cs1C1 | Cm | MS | 0.86+/-0.02 | 0.66+/-0.01 | 0.77 |
| SSU_28 | LD36Cs2C1 | Am | Cryo-EM | 0.57+/-0.05 | 0.47+/-0.02 | 0.84 |
| SSU_33 | - | Um | Cryo-EM | 0.11+/-0.11 | 0.04+/-0.04 | 0.40 |
| SSU_38 | LD36Cs2C1 | Cm | Cryo-EM | 0.40+/-0.01 | 0.34+/-0.00 | 0.83 |
| SSU_479 | LD32Cs1C1 | Am | Cryo-EM | 0.19+/-0.19 | 0.54+/-0.02 | 2.80 |
| SSU_509 | - | Gm | Cryo-EM | 0.00+/-0.00 | 0.12+/-0.03 | NA |
| SSU_512 | LD33Cs3C1 | Am | MS | 0.56+/-0.18 | 0.78+/-0.02 | 1.39 |
| SSU_661 | LD35Cs3C1/  LD35Cs3C-1 | Um | Cryo-EM | 0.36+/-0.21 | 0.75+/-0.03 | 2.06 |
| SSU_668 | LD35Cs3C1 | Am | Cryo-EM | 0.67+/-0.10 | 0.80+/-0.02 | 1.20 |
| SSU_8 | LD33Cs2C1 | Um | Cryo-EM | 0.60+/-0.07 | 0.67+/-0.00 | 1.12 |
| SSU_912 | LD34Cs1C2 | Am | Cryo-EM | 0.68+/-0.07 | 0.62+/-0.02 | 0.92 |
| SSU_98 | LD20Cs1C4 | Am | Cryo-EM | 0.67+/-0.13 | 0.82+/-0.02 | 1.22 |

*Footnote: Data are shown as mean ± s.e.m. The fold change (FC) of RMS of individual Nms is presented and the hypermodified sites showing FC > 2 are highlighted. Note that we have not identified all the snoRNAs guiding the existing Nm modification. (*) The rRNA unit (LSU, SSU or 5.8S) and the nucleotide carrying the Nm modification are reported.*
