## Supplementary material for "Genome instability drives epistatic adaptation in the human pathogen *Leishmania*": Table S21

**Table S21:** **The relative fold-change of Ψs in *L. donovani* rRNA between P135 and P2 parasites.**

| **rRNA** | **Position of Ψ** | **FC P135 vs P2 R1 (*)** | **FC P135 vs P2 R2 (*)** | **FC P135 vs P2 R3 (*)** | **FC P135 vs P2 R4 (*)** |
| --- | --- | --- | --- | --- | --- |
| SSU | 12 | 1.04 | 1.05 | 1.04 | 1.04 |
| SSU | 33 | 0.96 | 0.95 | 1.20 | 1.15 |
| SSU | 104 | 1.05 | 1.05 | 1.07 | 1.06 |
| SSU | 455 | 0.99 | 0.98 | 1.03 | 0.97 |
| SSU | 607 | 0.98 | 0.99 | 1.01 | 0.99 |
| SSU | 609 | 1.01 | 1.01 | 1.03 | 1.01 |
| SSU | 721 | 1.21 | 1.27 | 1.23 | 1.20 |
| SSU | 892 | 1.12 | 1.10 | 1.22 | 1.10 |
| SSU | 941 | 1.05 | 1.05 | 1.05 | 1.05 |
| SSU | 1156 | 1.05 | 1.06 | 1.06 | 1.03 |
| SSU | 1192 | 0.84 | 0.88 | 0.95 | 0.90 |
| SSU | 1246 | 1.02 | 1.02 | 1.05 | 1.03 |
| SSU | 1371 | 1.07 | 1.08 | 1.15 | 1.12 |
| SSU | 1374 | 1.21 | 1.29 | 1.26 | 1.35 |
| SSU | 1559 | 0.92 | 0.95 | 0.84 | 0.87 |
| SSU | 1566 | 0.97 | 0.98 | 0.97 | 0.97 |
| SSU | 1657 | 0.98 | 0.99 | 0.98 | 0.97 |
| SSU | 1722 | 1.02 | 1.02 | 1.07 | 1.04 |
| SSU | 1841 | 1.10 | 1.10 | 1.11 | 1.08 |
| SSU | 1970 | 1.10 | 1.09 | 1.18 | 1.05 |
| SSU | 2046 | 1.04 | 1.06 | 1.07 | 1.06 |
| SSU | 2048 | 1.09 | 1.09 | 1.10 | 1.10 |
| SSU | 2071 | 1.18 | 1.18 | 1.22 | 1.15 |
| 5.8S | 69 | 1.08 | 1.08 | 1.12 | 1.08 |
| 5.8S | 74 | 1.11 | 1.11 | 1.13 | 1.11 |
| LSUa | 239 | 1.06 | 1.06 | 1.12 | 1.03 |
| LSUa | 422 | 1.04 | 1.04 | 1.09 | 1.06 |
| LSUa | 672 | 1.14 | 1.11 | 1.15 | 1.09 |
| LSUa | 870 | 0.99 | 1.00 | 1.02 | 1.00 |
| LSUa | 1011 | 1.11 | 1.06 | 1.20 | 0.98 |
| LSUa | 1017 | 0.99 | 0.99 | 1.07 | 0.96 |
| LSUa | 1055 | 1.08 | 1.12 | 1.10 | 1.11 |
| LSUa | 1084 | 1.08 | 1.10 | 1.11 | 1.08 |
| LSUa | 1093 | 1.06 | 1.08 | 1.08 | 1.07 |
| LSUa | 1171 | 1.04 | 1.05 | 1.05 | 1.06 |
| LSUa | 1181 | 1.06 | 1.05 | 1.06 | 1.05 |
| LSUa | 1402 | 1.02 | 1.01 | 1.04 | 1.00 |
| LSUa | 1530 | 1.00 | 0.98 | 1.05 | 0.99 |
| LSUa | 1535 | 1.03 | 1.02 | 1.05 | 1.03 |
| LSUa | 1666 | 1.16 | 1.14 | 1.19 | 1.12 |
| LSUb | 78 | 0.97 | 0.99 | 1.03 | 1.01 |
| LSUb | 437 | 1.36 | 1.29 | 1.29 | 1.08 |
| LSUb | 472 | 1.00 | 1.25 | 1.17 | 1.37 |
| LSUb | 500 | 1.12 | 1.38 | 0.99 | 1.02 |
| LSUb | 504 | 1.15 | 1.54 | 1.26 | 1.30 |
| LSUb | 506 | 1.14 | 1.19 | 1.21 | 1.04 |
| LSUb | 510 | 1.11 | 1.16 | 1.17 | 1.09 |
| LSUb | 512 | 1.06 | 1.11 | 1.13 | 1.07 |
| LSUb | 593 | 1.03 | 1.45 | 1.01 | 1.21 |
| LSUb | 595 | 1.04 | 1.19 | 1.05 | 1.07 |
| LSUb | 597 | 1.02 | 1.15 | 1.04 | 1.06 |
| LSUb | 611 | 1.06 | 1.12 | 1.10 | 1.13 |
| LSUb | 626 | 1.02 | 1.04 | 1.04 | 1.05 |
| LSUb | 662 | 1.00 | 1.00 | 1.00 | 1.01 |
| LSUb | 704 | 1.00 | 0.99 | 1.00 | 1.00 |
| LSUb | 1059 | 1.08 | 1.07 | 1.14 | 1.07 |
| LSUb | 1061 | 1.04 | 1.04 | 1.08 | 1.05 |
| LSUb | 1144 | 0.51 | 0.69 | 0.32 | 1.45 |
| LSUb | 1265 | 1.43 | 2.85 | 1.49 | 2.18 |
| LSUb | 1282 | 1.35 | 2.25 | 1.30 | 1.85 |
| LSUb | 1285 | 1.34 | 1.79 | 1.47 | 1.43 |
| LSUb | 1304 | 1.32 | 1.79 | 1.38 | 1.50 |
| LSUb | 1319 | 1.30 | 1.64 | 1.41 | 1.50 |
| LSUb | 1355 | 1.27 | 1.62 | 1.36 | 1.41 |
| LSUb | 1362 | 1.18 | 1.39 | 1.21 | 1.28 |
| LSUb | 1383 | 1.10 | 1.18 | 1.14 | 1.14 |
| LSUb | 1404 | 1.08 | 1.09 | 1.10 | 1.09 |
| LSUb | 1414 | 1.08 | 1.09 | 1.09 | 1.09 |

*Footnote:(*) Four independent replicates (R). Only sites with log2 fold change (FC) >1.3 in all replicates are considered as hypermodified sites and are highlighted in pink.*
